# The recombination landscape of introgression in yeast

**DOI:** 10.1101/2024.01.04.574263

**Authors:** Enrique J. Schwarzkopf, Nathan Brandt, Caiti Smukowski Heil

**Affiliations:** Department of Biological Sciences, North Carolina State University, Raleigh, NC

**Keywords:** introgression, hybridization, recombination, crossover, non-crossover, yeast, Saccharomyces

## Abstract

Meiotic recombination is an evolutionary force that acts by breaking up genomic linkage, increasing the efficacy of selection. Recombination is initiated with a double-strand break which is resolved via a crossover, which involves the reciprocal exchange of genetic material between homologous chromosomes, or a non-crossover, which results in small tracts of non-reciprocal exchange of genetic material. Crossover and non-crossover rates vary between species, populations, individuals, and across the genome. In recent years, recombination rate has been associated with the distribution of ancestry derived from past interspecific hybridization (introgression) in a variety of species. We explore this interaction of recombination and introgression by sequencing spores and detecting crossovers and non-crossovers from two crosses of the yeast *Saccharomyces uvarum*. One cross is between strains which each contain introgression from their sister species, *S. eubayanus*, while the other cross has no introgression present. We find that the recombination landscape is significantly different between *S. uvarum* crosses, and that some of these differences can be explained by the presence of introgression in one cross. Crossovers are reduced and non-crossovers are increased in heterozygous introgression compared to syntenic regions in the cross without introgression. This translates to reduced allele shuffling within introgressed regions, and an overall reduction of shuffling on most chromosomes with introgression compared to the syntenic regions and chromosomes without introgression. Our results suggest that hybridization can significantly influence the recombination landscape, and that the reduction in allele shuffling contributes to the initial purging of introgression in the generations following a hybridization event.

## Introduction

Recombination is the exchange of genetic material between homologous chromosomes during meiosis and is a staple of eukaryotic sexual reproduction. While the processes involved in recombination are largely conserved (Arter & Keeney, 2023), recombination rates vary between sexes, populations, and species (Smukowski & Noor, 2011; Stapley et al., 2017).

Recombination rates also vary along the genome, with conflicting patterns of enriched or depleted recombination in promoter regions and punctate or dispersed recombination depending on the species (Auton et al., 2013; Rockman & Kruglyak, 2009; Singhal et al., 2015; Smukowski Heil et al., 2015). These patterns in recombination can affect pairing of alleles after meiosis–in other words, the shuffling of alleles–in a population. Much of the evolutionary advantage of recombination is understood to originate from its role in shuffling alleles, which increases the number of different allele combinations segregating in a population. The increase in allele combinations can reduce selection interference–the effect that genetically linked sites have on the evolutionary fate of either beneficial or deleterious alleles (Felsenstein, 1974; Hill & Robertson, 1966; McDonald et al., 2016; McGaugh et al., 2012).

How much allele decoupling is produced by recombination will depend on the type of recombination event. Each recombination event begins with the severing of both strands of a sister chromatid of one of the homologous chromosomes in what is referred to as a meiotic double-strand break (DSB) (Keeney, 2001). A meiotic DSB is repaired as a crossover (CO), which results in the reciprocal exchange of genetic information between homologous chromosomes, or as a non-crossover (NCO) gene conversion –where a small segment (typically 100-2000 bp) of a chromosome is replaced by a copy of its homolog (Chovnick et al., 1971; Hilliker et al., 1994; Jeffreys & May, 2004; Judd & Petes, 1988). COs generally produce more allele shuffling, and therefore degrade linkage faster than NCOs. However, NCOs can occur in regions where COs are typically suppressed, like centromeres and inversions (Korunes & Noor, 2019; Mancera et al., 2008; Miller et al., 2016; Schaeffer & Anderson, 2005; Shi et al., 2010; Talbert & Henikoff, 2010; Wijnker et al., 2013). NCOs are also crucial to reducing linkage within coding regions and, unlike COs, result in 3:1 allele ratio in the meiotic product at heterozygous sites, potentially changing allele frequencies (Korunes & Noor, 2017).

Variation in the number and distribution of COs and NCOs, and their respective associated effects on linkage, have important implications for molecular evolution. Recombination has long been appreciated to play a role in the distribution of various genomic features including nucleotide diversity. Nucleotide diversity has a positive correlation with recombination rate in a number of species, interpreted to result from selective sweeps and background selection removing genetic variation in regions of low recombination (Begun & Aquadro, 1992; Charlesworth et al., 1993; Smith & Haigh, 1974). Similarly, recombination breaking up genetic associations is particularly notable in the context of interspecific hybridization. In first-generation (F_1_) hybrids, the hybrid genome is heterozygous for each parental species. If the hybrids then back-cross to one of the parental species, recombination will produce genomes that are a mosaic of genetic information from the two species with a minor contribution from the species that was not backcrossed to (introgression) (Aguillon et al., 2022). When each population has evolved alleles that are deleterious when present in the background of the other population (the Dobzhansky-Muller hybrid incompatibility model) we expect introgressed regions with low rates of recombination to be quickly purged from the population, as the accumulation of incompatible alleles incurs a steep fitness cost. In contrast, when introgressed regions have high recombination rates, the break up of genetic associations will reduce selective interference between the incompatible alleles and their surrounding haplotypes, allowing for neutral and beneficial alleles brought in with the introgression to escape the fate of neighboring incompatibilities (Barton & Bengtsson, 1986; Butlin, 2005; Moran et al., 2021; Nachman, M. & Payseur, B., 2012; Schumer et al., 2018; Veller et al., 2023). This theory is supported empirically through enrichment of introgressed segments in regions of higher recombination in a number of organisms including Mimulus, maize, butterflies, swordtail fish, stickleback, and humans (Brandvain et al., 2014; Calfee et al., 2021; Edelman et al., 2019; Martin et al., 2019; Ravinet et al., 2018; Schumer et al., 2018).

This positive correlation between introgressed ancestry and recombination is emerging as a nearly ubiquitous pattern (though see (Dagilis & Matute, 2023; Duranton & Pool, 2022; Pool, 2015)), however, it is unclear how these observations relate to the known effect of sequence divergence on DSB repair. Introgression, particularly between highly diverged species, can have low sequence similarity with the genomic region it is replacing. A DSB in a region of low sequence similarity will recruit mismatch repair proteins, which ensure COs are occurring between homologous chromosomes and at equivalent positions to prevent ectopic recombination (Harfe & Jinks-Robertson, 2000; Hunter et al., 1996). Mismatch repair proteins typically reduce the frequency of CO events as sequence divergence increases (Borts & Haber, 1987; Chen & Jinks-Robertson, 1999; Cooper et al., 2021; Martini et al., 2011). Given that heterozygous introgression will have divergent sequences, we expect a decrease in COs, and possibly an increase in NCOs as DSBs fail to be repaired as COs in heterozygous introgression.

To help us understand this interaction of introgression and recombination, and identify patterns in CO and NCO in closely related populations, we utilized the budding yeast *Saccharomyces*. Yeasts provide an excellent opportunity to study DSB repair, as we can readily isolate and collect all four meiotic products of a given meiosis and detect both CO and more elusive NCO events (Figure 1A) (Brion et al., 2017; Gerton et al., 2000; Liu et al., 2018, 2019; Mancera et al., 2008). Recombination rates vary between strains of *S. cerevisiae* (Cubillos et al., 2011; Raffoux et al., 2018) and between *S. cerevisiae* and its sister species *S. paradoxus* (Liu et al., 2019; Tsai et al., 2010). Strains of different *Saccharomyces* species have often hybridized with other species and carry introgressed DNA from these events (Albertin et al., 2018; Almeida et al., 2014; Bendixsen et al., 2022; D’Angiolo et al., 2020; Langdon et al., 2019; Stelkens & Bendixsen, 2022; Tellini et al., 2023).

**Figure 1:**
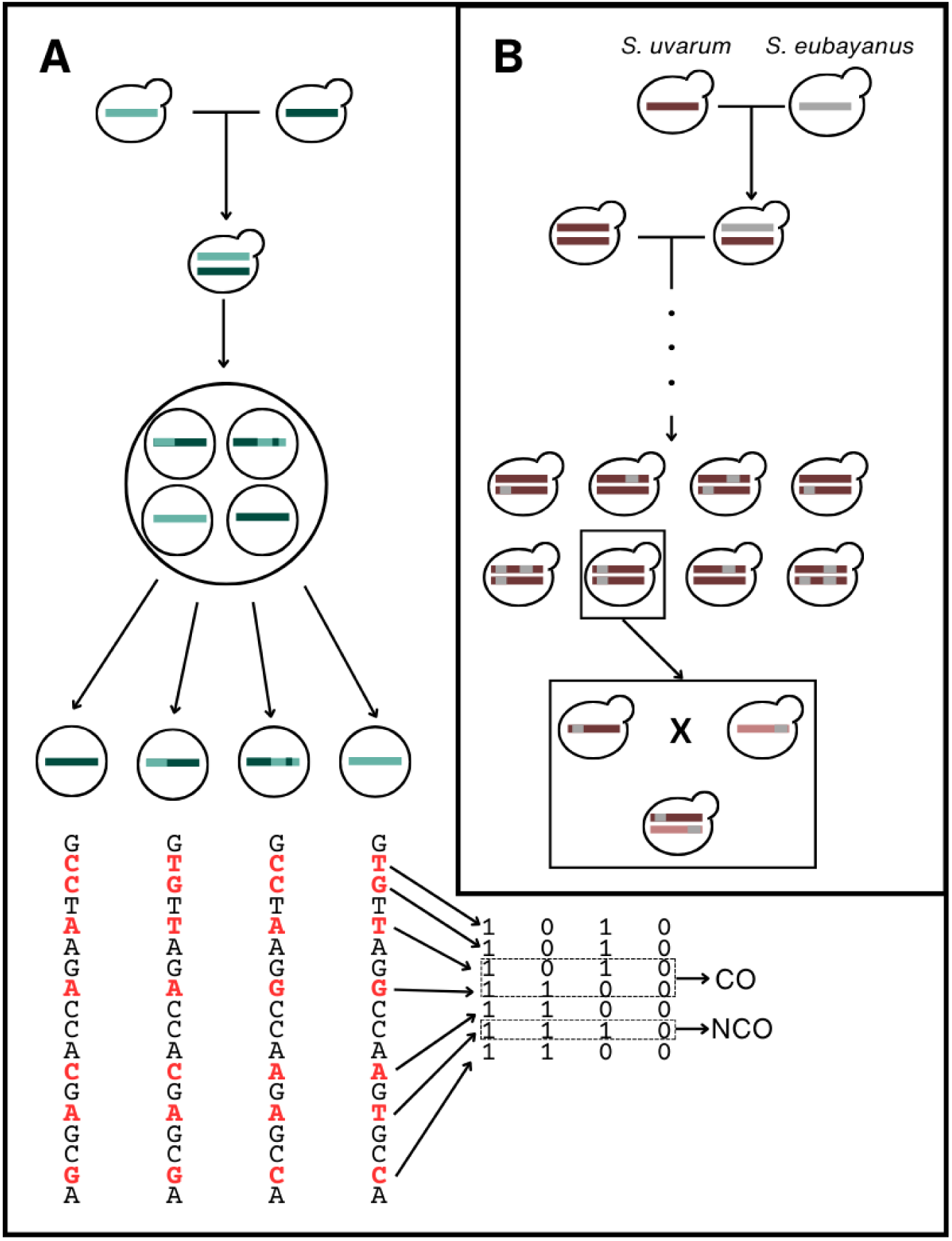
(A) Visual representation of how our crosses were conducted. Haploid yeast from two genetically distinct parental strains are mated to produce a heterozygous diploid. Meiosis is induced and the resulting meiotic products (tetrads) are manually dissected, and each haploid meiotic product is grown mitotically to obtain enough material for DNA extraction and whole genome sequencing. We then call SNPs on the resulting sequences and retain loci with fixed differences between parents. These loci are then coded as 1 or 0 depending on the parent of origin and the CrossOver software detects COs and NCOs. (B) A schematic of how introgression likely arose in the strains sampled from fermentation environments. These introgressions are likely due to S. eubayanus hybridizing with S. uvarum at some point in the past, resulting in F1 hybrids that then potentially crossed with other S. uvarum individuals for some number of generations. Eventually, the S. eubayanus ancestry was degraded in the population of S. uvarum until the introgressions we observe today remained, potentially segregating in the population. A similar process likely happened in each of the parental strains we utilized, but with different introgressions remaining in each strain. We crossed haploid individuals from two parental strains, resulting in a diploid that is heterozygous for each introgression.

In this study, we look at patterns of recombination and introgression by crossing two pairs of Holarctic *Saccharomyces uvarum* strains. One pair of strains was isolated from natural environments in North America and the other pair was isolated from European fermentation environments (Almeida et al., 2014). The *S. uvarum* strains isolated from European fermentation environments each carry introgression from their sister species, *Saccharomyces eubayanus*, which is approximately 6% divergent from *S. uvarum* (Almeida et al., 2014; Langdon et al., 2020; Nespolo et al., 2020). The diploid F_1_ genome of these strains is heterozygous for nine different introgressions which make up approximately 10% of the genome (Figure 1B). The strains from the North American cross do not carry *S. eubayanus* introgression, thus allowing us to assess the impact of introgression on the recombination landscape. We obtained whole genome sequencing data from individual meiotic events from the first offspring generation of each cross and used this data to detect CO and NCO events along the genome (Figure 1A). From these maps, we aim to understand (i) how patterns of CO and NCO differ between closely related strains, (ii) how regions of introgression differ in their CO and NCO patterns, and (iii) how these different patterns affect shuffling of alleles locally and at the chromosome level. Understanding these objectives will provide us novel insights into how introgression impacts the recombination landscape.

## Results

### The recombination landscape differs dramatically between closely related crosses

We isolated and sequenced products of 48 meioses (192 haploid spores) for two crosses of *S. uvarum*, a cross between strains isolated from North America (natural cross) and a cross between strains isolated from Europe (fermentation cross). We detected COs and NCOs across the 16 nuclear chromosomes of *S. uvarum*. Genomewide, we found significantly more COs on average in the natural cross (82.54 COs/meiosis, SE 1.5; 0.72 cM/kb) than in the fermentation cross (63.66 COs/meiosis, SE 1.9; 0.55 cM/kb; Wilcoxon rank sum test, p-value<<0.001) and significantly fewer NCOs in the natural cross (25.48 NCOs/meiosis, SE 0.99) than the fermentation cross (34.85 NCOs/meiosis, SE 8.92; Wilcoxon rank sum test, p-value=0.001448). The number of COs per meiosis in the natural and fermentation crosses are slightly higher than those of particular strains of *S. paradoxus* (54.8) and *S. cerevisiae* (76.5) respectively (Liu et al., 2019). Additionally, the average number of NCOs in the natural cross is lower than those of *S. cerevisiae* (46.4) and *S. paradoxus* (26.9), while the average number of NCOs in the fermentation cross is in between these values (Figure 2A; Tables S4 & S5; Liu et al., 2019). It’s important to note that there is known strain variability in CO and NCO counts per meiosis for *S. cerevisiae* (CO: 90.5, 76.3, 73; NCO: 46.6, 46.4, 27; Mancera et al., 2008; Liu et al., 2019; Martini et al., 2011), similar to differences between strains of *S. uvarum* reported here.

**Figure 2:**
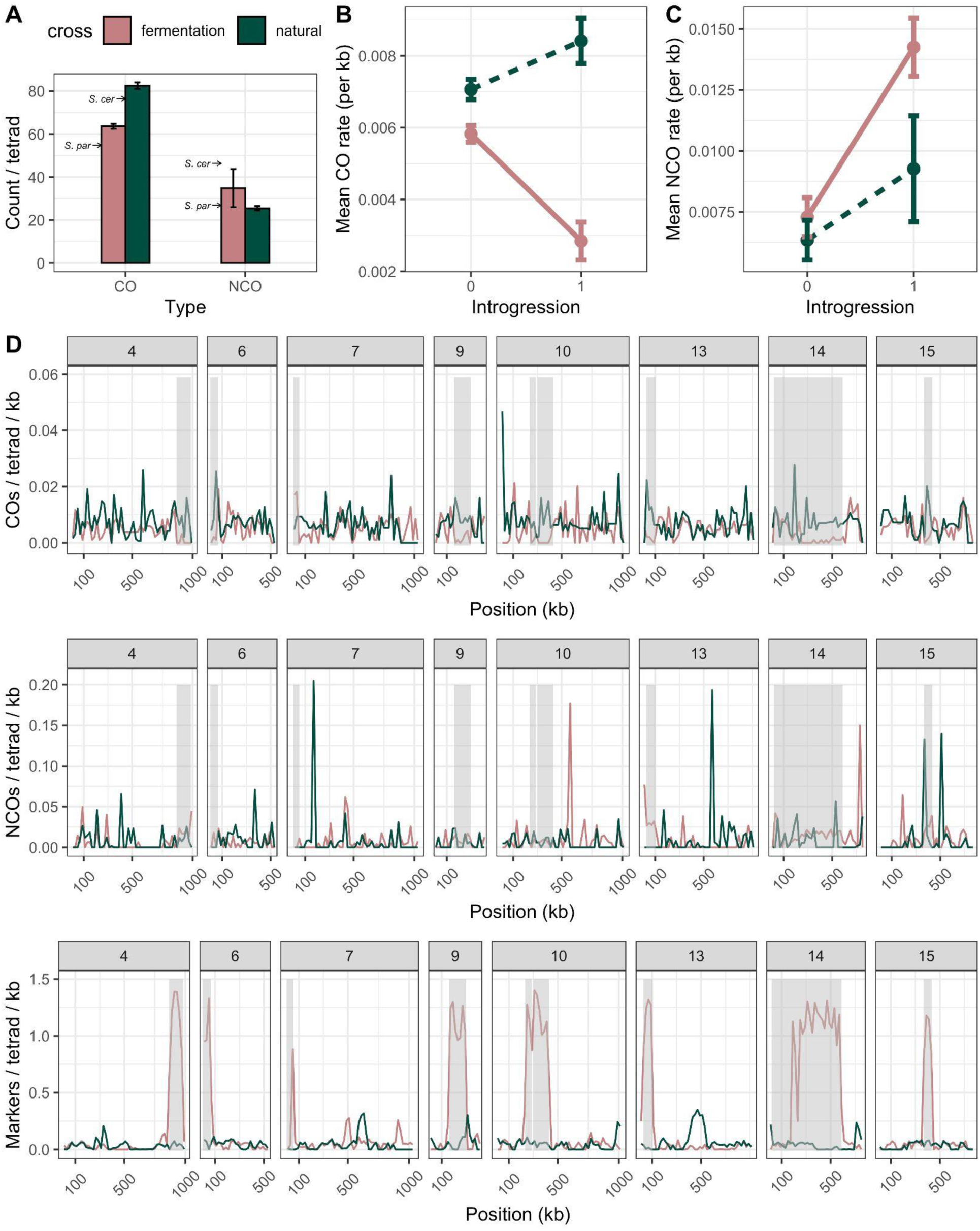
(A) Barplot depicting the number of COs and NCOs detected per meiosis in S. uvarum crosses (green: natural cross; pink: fermentation cross). The error bars represent the standard error around the mean. NCO counts are corrected for marker resolution.The counts for S. paradoxus and S. cerevisiae are represented by arrows and taken from Liu et al. (2019). (B) Mean CO/kb and (C) NCO/kb by cross and introgression (0 denotes intervals without introgression; 1 denotes introgression present in the fermentation cross. While the natural cross does not contain introgression, the region where introgression is present in the fermentation cross was compared to its syntenic region in the natural cross). NCO counts are corrected for marker resolution. Error bars represent the standard error around the mean. (D) S. uvarum chromosomes containing introgressions split into 20kb, non-overlapping windows. CO, NCO, and SNP counts are reported for both crosses (fermentation and natural). Shaded regions denote introgressed regions. CO counts are smoothed when the true location of the CO split could be in one of multiple windows. NCO counts are corrected for marker resolution.

To further explore the differences in recombination landscapes between our crosses, we split the genome into 20kb, non-overlapping windows, and obtained CO, NCO, and marker counts for each region. We downsampled markers in the introgression of the fermentation cross and found a significant effect of marker density on NCO count and tract length for most of the regions (Figures S2-10, Table S10). Because of how correlated marker density, NCO count, and introgression are, disentangling their relationships is complicated. For this reason, we applied an especially stringent correction for marker density–we used a previously published simulation based method for correction (Liu et al., 2019; Wijnker et al., 2013) and took the average NCO tract length (550bp) from introgression, where our marker density is highest, as the correction factor. For each window across the genome, we established our expected probability of detecting an NCO of that length and divided our observed NCO count by our probability of detecting an NCO event (see Methods). After the correction we found modest–but significant–genomewide correlation between our crosses for both COs (Spearman’s correlation: 0.27; p<0.0001) and NCOs (Spearman’s correlation: 0.13; p=0.0018).

We hypothesized that the low correlations between crosses might be impacted by the presence of heterozygous introgression from *S. eubayanus* in the fermentation cross. To explore this possibility, we separated the 20kb windows into introgressed and non-introgressed windows (based on whether they overlapped with an introgressed region). We will refer to introgression in the fermentation cross as “introgression” and use the term “introgressed region” to refer generally to the syntenic region, regardless of which cross we are focusing on. We calculated Spearman’s correlations of COs, NCOs, and marker counts between the two crosses for each chromosome. We find positive correlation coefficients between CO counts between the fermentation and natural crosses for all chromosomes when looking at non-introgressed regions–though only chromosome 11 was significant (Table S6). For introgressed regions, we found no significant CO correlations between crosses and no consistency in the direction of the correlations, which is consistent with the hypothesis that CO landscapes are changed in introgression (Table S7). There were no significant correlations among NCOs. This was likely affected by the fact that markers used to detect NCOs are differently distributed between the crosses, and the small size of NCO tracts means that the marker resolution of a given region will affect the reported location of NCOs. This is because NCOs are reported to be at the midpoint of the markers that identify it; more dense markers will give a more precise location, which may conflict with the location reported by less dense markers. We find that introgressions tend to have lower CO counts and higher NCO counts in the fermentation cross when compared to syntenic regions in the natural cross (Wilcoxon rank sum test; CO p-value<10^-5^, NCO p-value<10^5^). The fermentation cross has fewer COs than the natural cross overall, but the difference is greater in the introgressed regions (Figure 2B-C). Additionally, introgressed regions in the natural cross show significantly more COs and NCOs (Wilcoxon rank sum test; CO p-value=0.007, NCO p-value=0.006), indicating that these regions may be predisposed to having more recombination events even in the absence of introgression.

To further explore and test possible explanations for the patterns of COs and NCOs in introgressed regions, we constructed linear models for CO and NCO counts. Our model of CO counts showed a significant positive effect of the interactions between natural cross and introgression (whether a genomic window is from the natural cross and whether that window is in an introgressed region), as well as a significant negative effect of introgression on the number of COs, and a significant positive effect of GC content on CO count (Table 1). These results are consistent with increased COs in GC-rich regions and reduced COs in introgressions. Our model of NCO counts showed a similar positive effect of GC content on NCO counts, but showed an opposite significant coefficient for introgression and no significant effect of the interaction between introgression and cross–which was removed from the final model (Table 2). This indicates that GC content still plays an important role in localizing NCOs, and supports our findings that patterns of NCOs in introgressed regions are opposite to those of COs. Furthermore, the lack of a significant effect of cross echoes our finding of increased NCOs in introgressed regions in the natural cross.

**Table 1:** Coefficients of gaussian generalized linear model modeling CO counts per 20kb window.

|  | Estimate | Std. Error | t value | Pr(>\|t\|) |
| --- | --- | --- | --- | --- |
| (Intercept) | -8.9355 | 1.4713 | -6.0732 | 1.70e-09 |
| Introgression | -3.6343 | 0.6293 | -5.7755 | 9.84e-09 |
| GC | 37.9733 | 3.7037 | 10.2528 | < 2e-16 |
| Introgression:crossnatural | 5.2378 | 0.8595 | 6.0941 | 1.49e-09 |

**Table 2:** Coefficients of gaussian generalized linear model modeling NCO counts per 20kb window.

|  | Estimate | Std. Error | t value | Pr(>\|t\|) |
| --- | --- | --- | --- | --- |
| (Intercept) | -11.4763 | 4.7810 | -2.4004 | 0.0165 |
| Introgression | 4.3490 | 1.4937 | 2.9115 | 0.0037 |
| GC | 45.2894 | 12.0353 | 3.7631 | 0.0002 |

### Reduced diploid sequence similarity helps explain non-crossover repair of DSBs in introgression

One possible explanation for an increase of NCOs in introgression is that the reduced sequence similarity is biasing DSBs in the region to be repaired as NCOs rather than COs. We would therefore expect to see NCOs to be negatively correlated with sequence similarity. To evaluate the relationship of sequence similarity to the CO and NCO landscapes in introgressions, we measured diploid sequence similarity (the proportion of bases that are expected to match when we sample one base from each of the two parental strains), NCO depth, and CO count in 101 bp sliding windows with 50 bp overlaps along each of the introgressions. Mismatch repair proteins in *Saccharomyces* seem to suppress COs with very little mismatch in small regions (∼350 bp), which informed our window size (Chen & Jinks-Robertson, 1999). We counted the number of NCO tracts that intersect with each window as a measurement of NCO depth, and simply counted the CO events in a given window (Figure 3). We then ran Spearman’s correlations and a loess regression along each introgression and found a weak, but often significant (p<<0.001) correlation between NCOs and sequence similarity in the introgression (Table 3), suggesting that repair of double strand breaks is biased towards NCOs when sequence similarity is low. From the loess regression, we can observe an increase in NCO as sequence similarity reduces until about 0.9-0.8 sequence similarity, at which point NCOs level out or reduce (Figure S1). However, this effect is very weak with respect to the NCO counts, and at low levels of sequence similarity the uncertainty of the regression line is very large. This is primarily driven by the number of windows with no NCOs .

**Figure 3:**
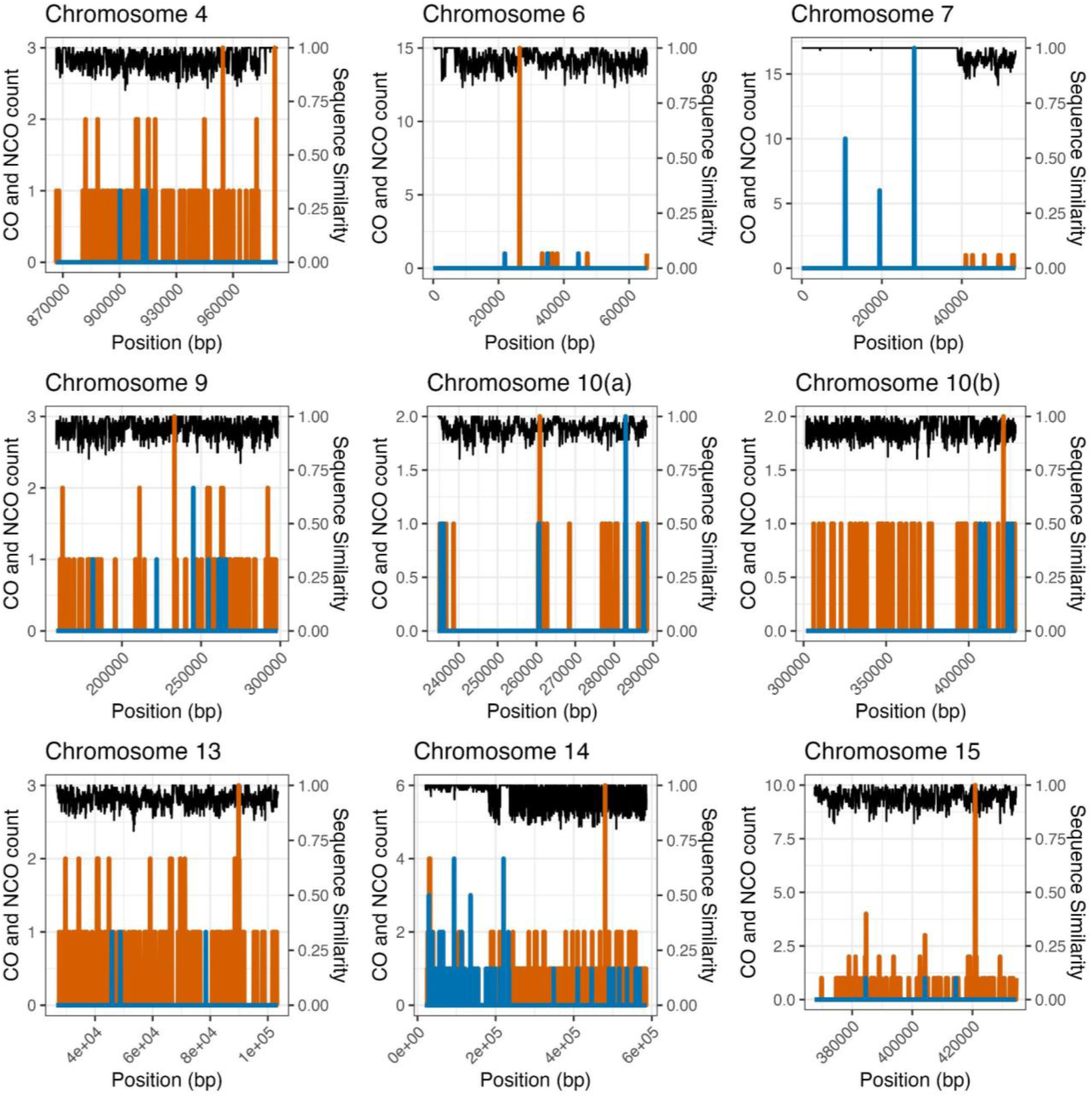
CO, NCO, and sequence similarity in 101bp sliding windows with 50bp overlaps of fermentation cross introgression. CO counts are shown in blue, the depth of NCO tracts are shown in orange, and the proportion of expected homologous bases between the two fermentation strains is shown in black.

**Table 3:** Spearman’s correlations of NCOs to sequence similarity in introgression in the fermentation cross. A cutoff p-value of 0.001 was selected by calculating the Bonferroni corrected *α*=0.05 for nine comparisons (0.0011) and rounding down.

| Chromosome | Start | End | Corr | p<0.001? |
| --- | --- | --- | --- | --- |
| 4 | 866500 | 983774 | -0.1686 | TRUE |
| 6 | 1 | 65500 | -0.0394 | FALSE |
| 7 | 1 | 53500 | -0.2435 | TRUE |
| 9 | 158500 | 298500 | -0.1460 | TRUE |
| 10 | 234500 | 288500 | 0.0073 | FALSE |
| 10 | 301500 | 428500 | -0.1097 | TRUE |
| 13 | 26500 | 103500 | -0.1982 | TRUE |
| 14 | 18500 | 586500 | -0.2300 | TRUE |
| 15 | 367500 | 434500 | -0.2328 | TRUE |

The low CO count in introgression leaves us unable to investigate effects of sequence similarity on CO counts except on chromosomes 7 and 14, where we found significantly higher sequence similarity around COs than around NCOs (Table 4). The introgressions on these two chromosomes are unique in that they contain a highly homologous portion of sequence and therefore contain enough COs for us to have power to detect differences between CO and NCO neighborhoods (Figure 3).

**Table 4:**
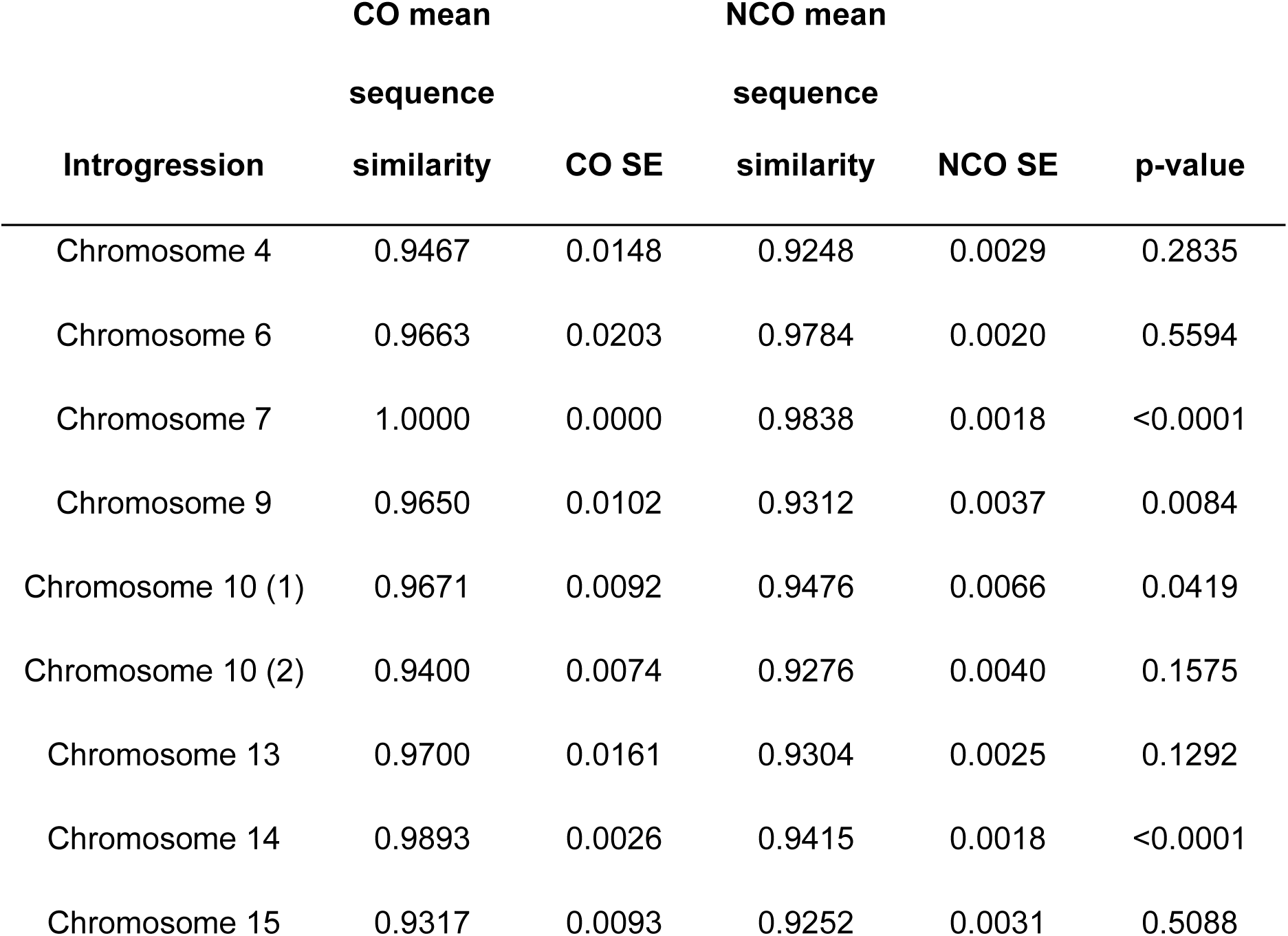
Welch two sample t-test results for differences in sequence similarity between CO-adjacent regions and NCO-adjacent regions per introgression.

### Introgression decreases allele shuffling locally and at the chromosome level

Because NCOs still play a small role in shuffling alleles along the chromosome, we were interested in whether the increase in NCOs of the fermentation cross would supplement the lost shuffling from the suppression of COs in the introgressions. To test this hypothesis, we used the measure *ṟ*, which accounts for the number and positioning of recombination events to estimate the probability that a randomly chosen pair of loci shuffles their alleles in a gamete (Veller et al., 2019). We calculated the average *ṟ* per chromosome and for each introgressed region for each of the two crosses. We observed high levels of shuffling at the chromosome level when compared to humans. The intra-chromosomal component of *ṟ* in humans is 0.0135 in females and 0.0177 in males (Veller et al., 2019), while our measurements for chromosomes varied between 0.216 and 0.426. We find that most chromosomes do not have a significantly different amount of allele shuffling between the two crosses, even though the natural cross generally has more COs (Table S8; Bonferroni-adjusted *α* = 0.00313). However, of the six chromosomes with significantly different *ṟ* values, all of them showed more shuffling in the natural cross, and five of the six (chromosomes 4, 9, 10, 14, and 15) contained introgressed regions (Figure 4). Lower *ṟ* is not observed when introgressions are small and near telomeres, while even a small introgression near the center of the chromosome can lead to a large reduction in *ṟ* (as is the case for chromosome 15). Chromosome 12 was the only chromosome without an introgressed region to have significantly different shuffling between crosses, and it also showed more shuffling in the natural cross. While it is unclear what potential mechanism is mediating the difference in shuffling on chromosome 12, we note that the rDNA locus on chromosome 12 is known to differ dramatically in repeat content across strains of *S. cerevisiae* (22–227 copies) (Sharma et al., 2022), and we speculate that differences in rDNA copy number between strains in our crosses could impact shuffling. All of the introgressed regions showed significantly more shuffling in the natural cross, indicating that the large increase of NCOs in the introgressions does not make up for the loss of shuffling from the depletion of COs (Table S9; Bonferroni-adjusted *α* = 0.00556). This finding indicates that an introgression that is segregating in a population will incur a shuffling cost in heterozygous individuals on top of any other evolutionary effects the introgression may have.

**Figure 4:**
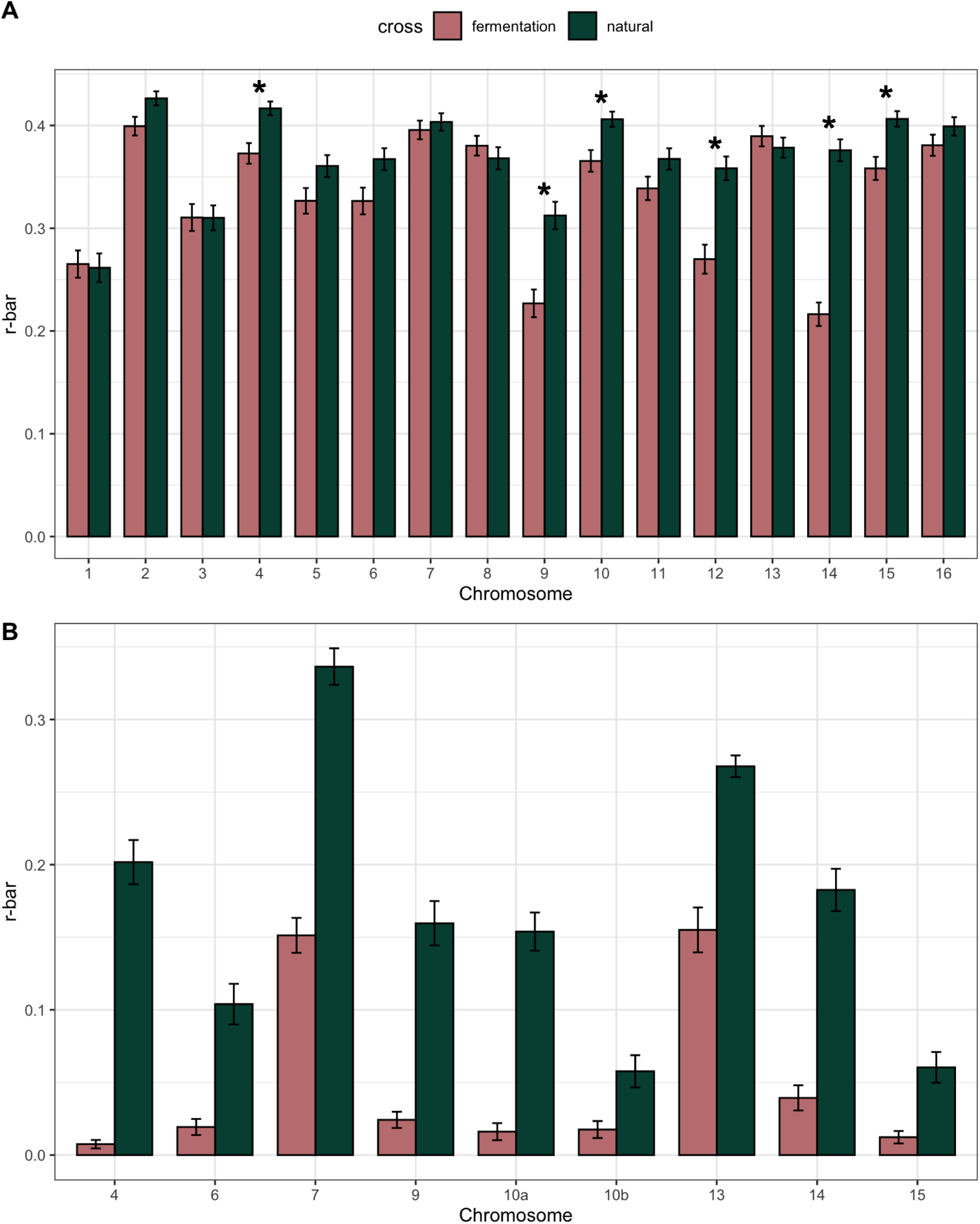
Average *ṟ* for each chromosome (A) and for each introgressed region (B). Asterisks indicate a significant difference in chromosome r-bar between crosses. All introgressed regions had a significant difference in *ṟ*. Error bars indicate standard error around the mean.

## Discussion

Our study is motivated by understanding recombination rate variation within a species and uncovering potential genetic factors underlying this variation. To investigate this question, we crossed two pairs of *S. uvarum* strains, one pair isolated from natural environments and one pair from fermentation environments, and explored the distribution of CO and NCO events from both crosses. We detected more COs and fewer NCOs in our natural cross when compared to our fermentation cross, within a similar range of COs and NCOs per meiosis as previous studies in *S. cerevisiae* and *S. paradoxus* (Liu et al., 2019; Mancera et al., 2008). Our findings demonstrate significant differences in recombination rate between closely related strains, adding to a body of literature across plants, animals, and fungi that recombination rate can evolve rapidly (Bauer et al., 2013; Danguy des Déserts et al., 2021; Kong et al., 2014; McGaugh et al., 2012; Petit et al., 2017; Raffoux et al., 2018; Samuk et al., 2019; Sandor et al., 2012; Schreiber et al., 2022).

We hypothesized that these differences in the recombination landscape between our crosses were in part influenced by introgression, given that heterozygous introgression creates sequence divergence, and that COs in regions of heterozygosity are known to be curtailed in yeast and other organisms (Chen & Jinks-Robertson, 1999; Cooper et al., 2021; Martini et al., 2011). We therefore explored the relationship between introgressions and the differences in CO and NCO counts between crosses. We modeled CO and NCO locations, correcting for GC content (a well characterized driver of recombination events (Kiktev et al., 2018; Marsolier-Kergoat & Yeramian, 2009)) and found that the distribution of COs and NCOs we observed could be partly explained by introgressions. While we are limited in our interpretations by only comparing two crosses (one cross with heterozygous introgression and one without introgression), these results are in line with findings in inversions, where heterozygotes show sharp decreases in COs, but the presence of NCOs in the inverted region (Crown et al., 2018; Korunes & Noor, 2019). However, unlike heterozygous inversions where an increase in COs is observed on freely recombining chromosomes (the inter-chromosomal effect), we do not see an increase in COs on the borders flanking introgression or on chromosomes without introgression. Relatedly, our results also differ from observations in Arabidopsis, in which COs are elevated in regions with more heterozygosity at the cost of COs in adjacent homologous regions (Ziolkowski et al., 2015; Ziolkowski & Henderson, 2017). These contrasting outcomes are likely in part due to the difference between intraspecific levels of polymorphism in Arabidopsis studies compared to the high level of inter-specific sequence divergence in our heterozygous introgression, although it also appears that mismatch repair gene MSH2 has functionally diverged between Arabidopsis and Saccharomyces (Blackwell et al., 2020; Dluzewska et al., 2023; Szymanska-Lejman et al., 2023). Regardless, it is curious that we do not find evidence for compensation of the depletion of COs due to introgression; the underlying mechanism responsible requires further study.

The introgressions present in the fermentation cross strains provided a unique opportunity to analyze NCO patterns, as the highly increased marker density lends the ability to detect NCOs with smaller tract lengths. We find evidence that DSB repair is biased towards NCOs when sequence similarity is low, with a weak but significant effect of increasing NCOs with decreasing sequencing similarity to a peak at sequence similarity between 0.8-0.9. Our data are thus aligned with previous work in *S. cerevisiae* and other systems, in which *MSH2* and *SGS1* act as anti-recombinases and disassemble D-loops with mismatched sequences, resulting in NCOs instead of COs (Borts & Haber, 1987; Chen & Jinks-Robertson, 1999; Martini et al., 2011; Mazina et al., 2004; Myung et al., 2001). By downsampling markers in the introgressed regions, we found an increase in the number of NCOs and a decrease in their tract length as we increased our resolution (Figures S2-S10). Our expectation was to find a marker density at which NCO counts and their tract lengths would no longer increase and decrease respectively. However, we didn’t find such a density, despite in some cases exploring marker densities that included all the markers in the introgression. This leaves us with the uncertainty of whether NCOs in the introgressions are more frequent and smaller than in other regions, or if we may be underestimating NCO counts and overestimating their tract lengths in cases where marker resolution is lower.

One likely effect of the CO reduction we observed in introgressions is a reduction in allele shuffling at the regional and chromosomal level. While NCOs can increase local shuffling, they likely have a much weaker effect on the likelihood of two random alleles being shuffled than COs do. We find this is the case for our two crosses, where despite a large number of NCOs in introgressions, the amount of shuffling (as measured by *ṟ*) is significantly lower in the fermentation cross. This loss of shuffling translates to frequently lower *ṟ* in the fermentation cross at the chromosome level for chromosomes containing introgressions. The exceptions being small introgressions near the telomeres, which is consistent with the expectation that COs near the center of chromosomes generate much more shuffling of alleles than terminal COs (Veller et al., 2019). Our findings indicate that reducing COs, especially near the center of chromosomes, has a cost to shuffling that is not compensated by the increase of NCOs that we observe. If the benefit of recombination is its ability to generate new combinations of alleles, then the loss of shuffling resulting from being heterozygous for divergent DNA sequences may come at an additional cost beyond the possibilities of genetic incompatibilities between hybridizing species. This cost is likely higher as divergence increases and as the length of divergent sequences is greater, as is the case with early generation hybrids (Dagilis & Matute, 2023). Ultimately, if sequence divergence is too high, the resultant failure to recombine can become a postzygotic reproductive barrier (Bozdag et al., 2021; Hunter et al., 1996; Rogers et al., 2018).

The shuffling cost to introgression that we identify in our crosses may play an important role in the fate of introgression in the generations following hybridization. When heterozygotes for an introgression are formed, the reduction in shuffling inside the introgression will increase the likelihood that the introgression is purged from the population. This is because it will likely be inherited in its entirety and will carry the fitness cost of incompatibilities combined with a cost of shuffling. This cost is incurred because the reduction of COs in the introgression will reduce shuffling of alleles on either side of it and will vary in its intensity depending on the location and size of the introgression. In generations immediately following hybridization, introgressions will be much larger and are therefore expected to be more costly (although this likely depends on a number of factors including time since divergence). These predictions are consistent with modeling and empirical data on the purging of introgression in Drosophila and humans in the first generations following hybridization (Veller et al., 2023).

As to longer term dynamics of recombination and selection, we predict that the excess NCOs detected in heterozygous introgressions should begin to erode the divergence between the sequences, increasing homology and slowly reducing the cost of the introgression. This hypothesis posits that recombination can act to remove the larger, more deleterious regions of an introgression quickly while whittling away slightly deleterious alleles that may be linked to any beneficial regions of an introgression. While our current study doesn’t capture longer term patterns of recombination or the landscape of recombination in introgressions that would lead to introgressions preferentially remaining in high-CO regions, it’s notable that introgressed regions coincide with high-recombination regions in the natural cross, indicating that these regions may be primed to retain introgressions after a hybridization event–possibly due to the increased shuffling allowing for a larger fraction of the introgression to remain (Veller et al., 2023).

Finally, we note that *Saccharomyces* typically reproduce asexually, with only infrequent sexual cycles (Magwene et al., 2011; Ruderfer et al., 2006; Zeyl & Otto, 2007). When they do mate, they often mate within a tetrad resulting in increased homozygosity. For example, each diploid progenitor of the parents of our fermentation cross was homozygous for introgression across the genome, meaning that recombination would neither break up nor aid in purging the introgression in isolated populations of each parent. This suggests that the fate of introgressions in this species is perhaps more loosely tied to recombination patterns than it would be in an obligately sexually reproducing species.

Despite some limitations to interpretation, this study provides a unique view of the early dynamics of hybridization and the role of recombination in the presence of introgression. By focusing not only on the distribution of recombination events but on their specific role in shuffling alleles, we can more closely connect the physical process of recombination to its role among other evolutionary forces.

## Methods

### Strain and library construction

*S. uvarum* strains (UCD61-137, yHCT78, GM14, and DBVPG7787) were obtained from the Portuguese Yeast Culture Collection and from Chris Hittinger (Table S1) (Almeida et al., 2014). All four *S. uvarum* strains had their *HO* locus replaced with a kanMX marker using a modified version of the high-efficiency yeast transformation using the LiAc/SS carrier DNA/PEG method. Briefly, the kanMX marker was amplified from plasmid pCSH2 with homology to genomic DNA flanking the HO ORF with primers CSH239 (GGTGGAAAACCACGAAAAGTTAGAACTACGTTCAGGCAAAgacatggaggcccagaatac) and CSH241 (GTGACCGTATTGGTACTTTTTTTGTTACCTGTTTTAGTAGcagtatagcgaccagcattc).

For each strain, overnight cultures were inoculated in 25 mL of YPD at an OD of ∼ 0.0005 and incubated at room temperature on a shaker for ∼24 hours until the cultures reached an OD between 0.6 and 0.9. Subsequently, 1 ug of the template DNA was transformed with a heat shock temperature of 37°C for 45 minutes. The transformed cells were allowed to recover in liquid YPD for 4 hours before being plated onto G418 selective plates and incubated at room temperature for 2 days.

Single colonies were selected from the transformation plates, restreaked onto G418 plates and allowed to grow at room temperature for 2 days. Single colonies from those plates were then inoculated into 2 mL of YPD + G418 and incubated in a roller drum at room temperature overnight. From those cultures, 250 uL was used to inoculate 2 mL of sporulation media (1% potassium acetate, 0.1 % yeast extract, 0.05% dextrose) and incubated at room temperature for 3 to 5 days. Strains were confirmed to have the ho::KanMX via tetrad dissection on a Singer SporPlay+ microscope (Singer Instruments). Plates with tetrads were incubated at room temperature for 2 days and then replica plated to test for proper segregation of the kanMX marker and mating type within individual tetrads.

Crosses between strains UCD61-137 and yHCT78 (natural cross), and between strains GM14 and DBVPG7787 (fermentation cross) were set up by micromanipulation of single MATa and MATx cells using a Singer SporPlay+. The plates were incubated at room temperature for 2 days and then replica plated to mating type tester strains to test for potential diploids. Identified diploids were then sporulated by growing a culture of the cross in 2 mL YPD + G418 at room temperature overnight. From those cultures, 250 uL were used to inoculate 2 mL of sporulation media and incubated at room temperature for 3 to 5 days. Sporulated cultures were dissected on 3 YPD plates (24 tetrads per plate) using a Singer SporPlay+. Fifty of the fully viable tetrads were selected and had all their spores inoculated into YPD (200 spores total) and incubated at room temperature. The DNA was extracted from these cultures using a modified version of the Hoffman-Winston DNA Prep (Hoffman & Winston, 1987). The DNA concentration was then measured using SYBR green, and 150 ng of each sample’s DNA was used to prepare a sequencing library using an Illumina DNA Prep Kit, modified to use half the normal amounts of reagents. Libraries were pooled and run on an Illumina NovaSeq 500 with 150bp paired end reads.

### Calling SNPs

We scored SNPs from parents and offspring using the *S. uvarum* reference genome (Scannell et al., 2011) and custom scripts that invoked bwa (v0.7.17), samtools (v1.12), bcftools (v1.13), picard tools (v2.25.6), and gatk (v4.2.0.0) (Danecek et al., 2021; Li & Durbin, 2009; McKenna et al., 2010). The custom scripts are available in the github repository: *schwarzkopf/CO-NCO.* We joint genotyped parents and offspring with default filters for gatk with the exception of the QUAL filter, which was set as < 100 for parents and < 30 for offspring. We further filtered variants by requiring they be fixed differences between the two parental strains. We kept a total of 24,574 markers for the natural cross and 74,619 markers for the fermentation cross. We utilized LUMPY to identify structural variants in the parent strains that were greater than 5000 bp and verified calls using the Integrative Genomics Viewer (Layer et al., 2014; Robinson et al., 2011). We identified three amplifications in strain GM14 (one of the fermentation cross parents) that were absent in other strains (Table S2).

### Generating CO/NCO maps

We generated “seg” files by coding tetrad variants by their parental origin. These seg files were the input for CrossOver (v6.3) from the ReCombine suite of programs, which we used to detect COs and NCOs (Anderson et al., 2011). We then filtered to remove non-crossovers with fewer than three associated markers and split the genome into 20kb windows. In each window we counted crossovers, non-crossovers, and markers. We established regions of introgressions through visual inspection of marker density in the fermentation cross (introgressions showed more divergence between fermentation strains) and confirmed them using the findings of Almeida et al. (2014). We found nine heterozygous introgressions on chromosomes 4, 6, 7, 9, 10, 10, 13, 14, and 15 respectively that we included in further analyses (Table S3). We excluded two additional introgressions due to poor mapping (chromosome 13:0-17,000; chromosome 16: 642,000-648,000). We analyzed the effect of marker resolution on NCO detection by randomly removing markers from each introgression and running CrossOver and its downstream counting process as described above. For each introgression, we downsampled to multiple percentiles of the distribution of non-introgressed region marker counts per kb (10^th^: 0.05; 25^th^: 0.75; 50^th^: 1.95; 75^th^: 2.95; 90^th^: 4.45; 95^th^: 7.87; 99^th^: 14.01) and the median marker count per kb for introgressed (3.225) and non-introgressed (1.35) regions. We repeated each downsampling amount 20 times for each introgression and extracted average CO counts, NCO counts, and NCO tract lengths. To account for the difference in number of markers in introgressed vs non-introgressed windows and their effect on NCO detection, we applied a previously published simulation-based method (Liu et al., 2019; Wijnker et al., 2013). We chose the average NCO tract length of the fermentation cross’s introgressed regions–550 bp–and for each window randomly placed an NCO event of that length 10,000 times to establish our expected probability of detecting an NCO of that length. We then divided our observed NCO count by our probability of detecting an NCO event. Additionally, because COs that occurred in large regions devoid of markers would be called in the middle of the empty windows, we split CO counts in regions with multiple consecutive marker-free windows evenly between the empty windows. With these corrected maps, we calculated spearman correlations between crosses using R (v4.1.0, R Core Team 2021). Additionally, we modeled NCO and CO count as a function of introgression, introgression by cross, and GC content using a gaussian generalized linear model in R (v4.1.0, R Core Team 2021).

### Sequence similarity

We calculated diploid sequence similarity between the two fermentation cross strains in 101bp windows with 50bp overlaps. At each nucleotide position in the window, we counted fixed differences as zero diploid sequence similarity, invariant sites between strains as full diploid sequence similarity (1), and polymorphic sites in either or both strains as half diploid sequence similarity (0.5). We then averaged these sequence similarity values across the window. This measure represents the probability that both strains will have the same nucleotide base at a given position. We used this measure of fine-scale sequence similarity to determine how sequence similarity related to NCO counts in introgressed regions. For this, we used Loess regressions and Spearman’s correlations on each of the introgressed regions comparing sequence similarity to NCO count, both implemented in R (v4.1.0, R Core Team 2021). We then focused on each recombination event (CO or NCO) and compared the sequence similarity 100bp up and downstream of CO breakpoints and 100bp up and downstream of NCO tracts.

We then used Welch’s two sample t-tests to compare CO and NCO sequence similarity in each introgression.

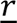

We use *ṟ*, a measure genetic shuffling defined in Veller et al. (2019) to measure how much shuffling occurs in each chromosome for each cross. Our data provides parental origin for each fixed difference between parental strains. We assume that all loci between pairs of markers that come from the same parent are also from that parent. We also assume that when a pair of successive markers come from different parents, the location of the change from one parental origin to the other happens at the midpoint between our markers. With this in mind, we counted the number of bases that come from one parent and divided by the chromosome size to obtain the proportion of the chromosome that was inherited from said parent (*p*) and used the formula from Veller et al. (2019): *ṟ* = 2*p*(1 − *p*). We calculated *ṟ* for each full chromosome and each introgressed regions in every gamete from both crosses. We then averaged across gametes to obtain average *ṟ* values. We then compared average *ṟ* between crosses in each chromosome or introgressed region using Welch two sample t-tests and correcting for multiple tests using a Bonferroni correction in R (v4.1.0, R Core Team 2021).

## Data Access

Sequences for the parental strains can be found on NCBI SRA (SRR1119189, SRR1119180 SRR1119199, SRR1119200) (Almeida et al., 2014). Sequencing of the tetrads is deposited at NCBI SRA under Project PRJNA1061120. Scripts are available in the github repository: *ejschwarzkopf/CO-NCO*.

## Acknowledgements

We are grateful to members of the Heil lab, Mohamed Noor, Nathan Layman, Mark Smithson, and three anonymous reviewers for comments on this manuscript. We thank Chris Hittinger and the Portuguese Yeast Culture Collection for *S. uvarum* strains. This work was supported by NIH R35GM142849 to CSH.

## References

1. Blackwell, Alexander & Dluzewska, Julia & Szymanska-Lejman, Maja & Desjardins, Stuart & Tock, Andrew & Kbiri, Nadia & Lambing, Christophe & Lawrence, Emma & Bieluszewski, Tomasz & Rowan, Beth & Higgins, James & Ziolkowski, Piotr & Henderson, Ian. (2020). MSH2 shapes the meiotic crossover landscape in relation to interhomolog polymorphism in Arabidopsis. The EMBO journal. 39. 10.15252/embj.2020104858.

2. Martini, Emmanuelle & Borde, Valérie & Legendre, Matthieu & Audic, Stéphane & Regnault, Beatrice & Soubigou, Guillaume & Dujon, Bernard & Llorente, Bertrand. (2011). Genome-Wide Analysis of Heteroduplex DNA in Mismatch Repair–Deficient Yeast Cells Reveals Novel Properties of Meiotic Recombination Pathways. PLoS genetics. 7. e1002305. 10.1371/journal.pgen.1002305.

3. Aguillon, S. M., Dodge, T. O., Preising, G. A., & Schumer, M. (2022). Introgression. Current Biology, 32(16), R865–R868. 10.1016/j.cub.2022.07.004

4. Albertin, W., Chernova, M., Durrens, P., Guichoux, E., Sherman, D. J., Masneuf-Pomarede, I., & Marullo, P. (2018). Many interspecific chromosomal introgressions are highly prevalent in Holarctic Saccharomyces uvarum strains found in human-related fermentations. Yeast, 35(1), 141–156. 10.1002/yea.3248

5. Almeida, P., Gonçalves, C., Teixeira, S., Libkind, D., Bontrager, M., Masneuf-Pomarède, I., Albertin, W., Durrens, P., Sherman, D. J., Marullo, P., Todd Hittinger, C., Gonçalves, P., & Sampaio, J. P. (2014). A Gondwanan imprint on global diversity and domestication of wine and cider yeast *Saccharomyces uvarum*. Nature Communications, 5, 4044. 10.1038/ncomms5044

6. Anderson, C. M., Chen, S. Y., Dimon, M. T., Oke, A., DeRisi, J. L., & Fung, J. C. (2011). ReCombine: A Suite of Programs for Detection and Analysis of Meiotic Recombination in Whole-Genome Datasets. PLoS ONE, 6(10). 10.1371/journal.pone.0025509

7. Arter, M., & Keeney, S. (2023). Divergence and conservation of the meiotic recombination machinery. Nature Reviews Genetics, 1–17. 10.1038/s41576-023-00669-8

8. Auton, A., Li, Y. R., Kidd, J., Oliveira, K., Nadel, J., Holloway, J. K., Hayward, J. J., Cohen, P. E., Greally, J. M., Wang, J., Bustamante, C. D., & Boyko, A. R. (2013). Genetic Recombination Is Targeted towards Gene Promoter Regions in Dogs. PLOS Genetics, 9(12), e1003984. 10.1371/journal.pgen.1003984

9. Barton, N., & Bengtsson, B. O. (1986). The barrier to genetic exchange between hybridising populations. Heredity, 57(3), Article 3. 10.1038/hdy.1986.135

10. Bauer, E., Falque, M., Walter, H., Bauland, C., Camisan, C., Campo, L., Meyer, N., Ranc, N., Rincent, R., Schipprack, W., Altmann, T., Flament, P., Melchinger, A. E., Menz, M., Moreno-González, J., Ouzunova, M., Revilla, P., Charcosset, A., Martin, O. C., & Schön, C.-C. (2013). Intraspecific variation of recombination rate in maize. Genome Biology, 14(9), R103. 10.1186/gb-2013-14-9-r103

11. Begun, D. J., & Aquadro, C. F. (1992). Levels of naturally occurring DNA polymorphism correlate with recombination rates in D. melanogaster. Nature, 356(6369), 519–520. 10.1038/356519a0

12. Bendixsen, D. P., Frazão, J. G., & Stelkens, R. (2022). Saccharomyces yeast hybrids on the rise. Yeast, 39(1–2), 40–54. 10.1002/yea.3684

13. Blackwell, A. R., Dluzewska, J., Szymanska-Lejman, M., Desjardins, S., Tock, A. J., Kbiri, N., Lambing, C., Lawrence, E. J., Bieluszewski, T., Rowan, B., Higgins, J. D., Ziolkowski, P. A., & Henderson, I. R. (2020). MSH2 shapes the meiotic crossover landscape in relation to interhomolog polymorphism in Arabidopsis. The EMBO Journal, 39(21), e104858. 10.15252/embj.2020104858

14. Borts, R. H., & Haber, J. E. (1987). Meiotic Recombination in Yeast: Alteration by Multiple Heterozygosities. Science, 237(4821), 1459–1465. 10.1126/science.2820060

15. Bozdag, G. O., Ono, J., Denton, J. A., Karakoc, E., Hunter, N., Leu, J.-Y., & Greig, D. (2021). Breaking a species barrier by enabling hybrid recombination. Current Biology, 31(4), R180–R181. 10.1016/j.cub.2020.12.038

16. Brandvain, Y., Kenney, A. M., Flagel, L., Coop, G., & Sweigart, A. L. (2014). Speciation and Introgression between Mimulus nasutus and Mimulus guttatus. PLOS Genetics, 10(6), e1004410. 10.1371/journal.pgen.1004410

17. Brion, C., Legrand, S., Peter, J., Caradec, C., Pflieger, D., Hou, J., Friedrich, A., Llorente, B., & Schacherer, J. (2017). Variation of the meiotic recombination landscape and properties over a broad evolutionary distance in yeasts. PLOS Genetics, 13(8), e1006917. 10.1371/journal.pgen.1006917

18. Butlin, R. K. (2005). Recombination and speciation. Molecular Ecology, 14(9), 2621–2635. 10.1111/j.1365-294X.2005.02617.x

19. Calfee, E., Gates, D., Lorant, A., Perkins, M. T., Coop, G., & Ross-Ibarra, J. (2021). Selective sorting of ancestral introgression in maize and teosinte along an elevational cline. bioRxiv, 2021.03.05.434040. 10.1101/2021.03.05.434040

20. Charlesworth, B., Morgan, M. T., & Charlesworth, D. (1993). The effect of deleterious mutations on neutral molecular variation. Genetics, 134(4), 1289–1303. 10.1093/genetics/134.4.1289

21. Chen, W., & Jinks-Robertson, S. (1999). The role of the mismatch repair machinery in regulating mitotic and meiotic recombination between diverged sequences in yeast. Genetics, 151(4), 1299–1313.

22. Chovnick, A., Ballantyne, G. H., & Holm, D. G. (1971). Studies on gene conversion and its relationship to linked exchange in Drosophila melanogaster. Genetics, 69(2), 179–209. 10.1093/genetics/69.2.179

23. Cooper, T. J., Crawford, M. R., Hunt, L. J., Marsolier-Kergoat, M.-C., Llorente, B., & Neale, M. J. (2021). Mismatch repair disturbs meiotic class I crossover control (p. 480418). bioRxiv. 10.1101/480418

24. Crown, K. N., Miller, D. E., Sekelsky, J., & Hawley, R. S. (2018). Local Inversion Heterozygosity Alters Recombination throughout the Genome. Current Biology: CB, 28(18), 2984–2990.e3. 10.1016/j.cub.2018.07.004

25. Cubillos, F. A., Billi, E., Zörgö, E., Parts, L., Fargier, P., Omholt, S., Blomberg, A., Warringer, J., Louis, E. J., & Liti, G. (2011). Assessing the complex architecture of polygenic traits in diverged yeast populations. Molecular Ecology, 20(7), 1401–1413. 10.1111/j.1365-294X.2011.05005.x

26. Dagilis, A. J., & Matute, D. R. (2023). The fitness of an introgressing haplotype changes over the course of divergence and depends on its size and genomic location. PLOS Biology, 21(7), e3002185. 10.1371/journal.pbio.3002185

27. Danecek, P., Bonfield, J. K., Liddle, J., Marshall, J., Ohan, V., Pollard, M. O., Whitwham, A., Keane, T., McCarthy, S. A., Davies, R. M., & Li, H. (2021). Twelve years of SAMtools and BCFtools. GigaScience, 10(2), giab008. 10.1093/gigascience/giab008

28. D’Angiolo, M., De Chiara, M., Yue, J.-X., Irizar, A., Stenberg, S., Persson, K., Llored, A., Barré, B., Schacherer, J., Marangoni, R., Gilson, E., Warringer, J., & Liti, G. (2020). A yeast living ancestor reveals the origin of genomic introgressions. Nature, 587(7834), Article 7834. 10.1038/s41586-020-2889-1

29. Danguy des Déserts, A., Bouchet, S., Sourdille, P., & Servin, B. (2021). Evolution of Recombination Landscapes in Diverging Populations of Bread Wheat. Genome Biology and Evolution, 13(8), evab152. 10.1093/gbe/evab152

30. Dluzewska, J., Dziegielewski, W., Szymanska-Lejman, M., Gazecka, M., Henderson, I. R., Higgins, J. D., & Ziolkowski, P. A. (2023). MSH2 stimulates interfering and inhibits non-interfering crossovers in response to genetic polymorphism. Nature Communications, 14, 6716. 10.1038/s41467-023-42511-z

31. Duranton, M., & Pool, J. E. (2022). Interactions Between Natural Selection and Recombination Shape the Genomic Landscape of Introgression. Molecular Biology and Evolution, 39(7), msac122. 10.1093/molbev/msac122

32. Edelman, N. B., Frandsen, P. B., Miyagi, M., Clavijo, B., Davey, J., Dikow, R. B., García-Accinelli, G., Van Belleghem, S. M., Patterson, N., Neafsey, D. E., Challis, R., Kumar, S., Moreira, G. R. P., Salazar, C., Chouteau, M., Counterman, B. A., Papa, R., Blaxter, M., Reed, R. D., … Mallet, J. (2019). Genomic architecture and introgression shape a butterfly radiation. Science, 366(6465), 594–599. 10.1126/science.aaw2090

33. Felsenstein, J. (1974). The evolutionary advantage of recombination. Genetics, 78(2), 737–756.

34. Gerton, J. L., DeRisi, J., Shroff, R., Lichten, M., Brown, P. O., & Petes, T. D. (2000). Global mapping of meiotic recombination hotspots and coldspots in the yeast Saccharomyces cerevisiae. Proceedings of the National Academy of Sciences, 97(21), 11383–11390. 10.1073/pnas.97.21.11383

35. Harfe, B. D., & Jinks-Robertson, S. (2000). Dna Mismatch Repair and Genetic Instability. Annual Review of Genetics, 34(1), 359–399. 10.1146/annurev.genet.34.1.359

36. Hill, W. G., & Robertson, A. (1966). The effect of linkage on limits to artificial selection. Genetical Research, 8(3), 269–294.

37. Hilliker, A. J., Harauz, G., Reaume, A. G., Gray, M., Clark, S. H., & Chovnick, A. (1994). Meiotic gene conversion tract length distribution within the rosy locus of Drosophila melanogaster. Genetics, 137(4), 1019–1026. 10.1093/genetics/137.4.1019

38. Hoffman, C. S., & Winston, F. (1987). A ten-minute DNA preparation from yeast efficiently releases autonomous plasmids for transformation of Escherichia coli. Gene, 57(2–3), 267–272. 10.1016/0378-1119(87)90131-4

39. Hunter, N., Chambers, S. R., Louis, E. J., & Borts, R. H. (1996). The mismatch repair system contributes to meiotic sterility in an interspecific yeast hybrid. The EMBO Journal, 15(7), 1726–1733. 10.1002/j.1460-2075.1996.tb00518.x

40. Jeffreys, A. J., & May, C. A. (2004). Intense and highly localized gene conversion activity in human meiotic crossover hot spots. Nature Genetics, 36(2), Article 2. 10.1038/ng1287

41. Judd, S. R., & Petes, T. D. (1988). Physical Lengths of Meiotic and Mitotic Gene Conversion Tracts in Saccharomyces Cerevisiae. Genetics, 118(3), 401. 10.1093/genetics/118.3.401

42. Keeney, S. (2001). Mechanism and control of meiotic recombination initiation. Current Topics in Developmental Biology, 52, 1–53. 10.1016/s0070-2153(01)52008-6

43. Kiktev, D. A., Sheng, Z., Lobachev, K. S., & Petes, T. D. (2018). GC content elevates mutation and recombination rates in the yeast *Saccharomyces cerevisiae*. Proceedings of the National Academy of Sciences, 115(30), E7109–E7118. 10.1073/pnas.1807334115

44. Kong, A., Thorleifsson, G., Frigge, M. L., Masson, G., Gudbjartsson, D. F., Villemoes, R., Magnusdottir, E., Olafsdottir, S. B., Thorsteinsdottir, U., & Stefansson, K. (2014). Common and low-frequency variants associated with genome-wide recombination rate. Nature Genetics, 46(1), 11–16. 10.1038/ng.2833

45. Korunes, K. L., & Noor, M. A. F. (2017). Gene conversion and linkage: Effects on genome evolution and speciation. Molecular Ecology, 26(1), 351–364. 10.1111/mec.13736

46. Korunes, K. L., & Noor, M. A. F. (2019). Pervasive gene conversion in chromosomal inversion heterozygotes. Molecular Ecology, 28(6), 1302–1315. 10.1111/mec.14921

47. Langdon, Q. K., Peris, D., Baker, E. P., Opulente, D. A., Nguyen, H.-V., Bond, U., Gonçalves, P., Sampaio, J. P., Libkind, D., & Hittinger, C. T. (2019). Fermentation innovation through complex hybridization of wild and domesticated yeasts. Nature Ecology & Evolution, 1–11. 10.1038/s41559-019-0998-8

48. Langdon, Q. K., Peris, D., Eizaguirre, J. I., Opulente, D. A., Buh, K. V., Sylvester, K., Jarzyna, M., Rodríguez, M. E., Lopes, C. A., Libkind, D., & Hittinger, C. T. (2020). Postglacial migration shaped the genomic diversity and global distribution of the wild ancestor of lager-brewing hybrids. PLOS Genetics, 16(4), e1008680. 10.1371/journal.pgen.1008680

49. Layer, R. M., Chiang, C., Quinlan, A. R., & Hall, I. M. (2014). LUMPY: A probabilistic framework for structural variant discovery. Genome Biology, 15(6), R84. 10.1186/gb-2014-15-6-r84

50. Li, H., & Durbin, R. (2009). Fast and accurate short read alignment with Burrows-Wheeler transform. *Bioinformatics (Oxford*, England), 25(14), 1754–1760. 10.1093/bioinformatics/btp324

51. Liu, H., Huang, J., Sun, X., Li, J., Hu, Y., Yu, L., Liti, G., Tian, D., Hurst, L. D., & Yang, S. (2018). Tetrad analysis in plants and fungi finds large differences in gene conversion rates but no GC bias. Nature Ecology & Evolution, 2(1), Article 1. 10.1038/s41559-017-0372-7

52. Liu, H., Maclean, C. J., & Zhang, J. (2019). Evolution of the Yeast Recombination Landscape. Molecular Biology and Evolution, 36(2), 412–422. 10.1093/molbev/msy233

53. Magwene, P. M., Kayıkçı, Ö., Granek, J. A., Reininga, J. M., Scholl, Z., & Murray, D. (2011). Outcrossing, mitotic recombination, and life-history trade-offs shape genome evolution in Saccharomyces cerevisiae. Proceedings of the National Academy of Sciences, 108(5), 1987–1992. 10.1073/pnas.1012544108

54. Mancera, E., Bourgon, R., Brozzi, A., Huber, W., & Steinmetz, L. M. (2008). High-resolution mapping of meiotic crossovers and non-crossovers in yeast. Nature, 454(7203), 479–485. 10.1038/nature07135

55. Marsolier-Kergoat, M.-C., & Yeramian, E. (2009). GC Content and Recombination: Reassessing the Causal Effects for the Saccharomyces cerevisiae Genome. Genetics, 183(1), 31–38. 10.1534/genetics.109.105049

56. Martin, S. H., Davey, J. W., Salazar, C., & Jiggins, C. D. (2019). Recombination rate variation shapes barriers to introgression across butterfly genomes. PLOS Biology, 17(2), e2006288. 10.1371/journal.pbio.2006288

57. Martini, E., Borde, V., Legendre, M., Audic, S., Regnault, B., Soubigou, G., Dujon, B., & Llorente, B. (2011). Genome-Wide Analysis of Heteroduplex DNA in Mismatch Repair– Deficient Yeast Cells Reveals Novel Properties of Meiotic Recombination Pathways. PLOS Genetics, 7(9), e1002305. 10.1371/journal.pgen.1002305

58. Mazina, O. M., Mazin, A. V., Nakagawa, T., Kolodner, R. D., & Kowalczykowski, S. C. (2004). Saccharomyces cerevisiae Mer3 helicase stimulates 3’-5’ heteroduplex extension by Rad51; implications for crossover control in meiotic recombination. Cell, 117(1), 47–56. 10.1016/s0092-8674(04)00294-6

59. McDonald, M. J., Rice, D. P., & Desai, M. M. (2016). Sex speeds adaptation by altering the dynamics of molecular evolution. Nature, 531(7593), 233–236. 10.1038/nature17143

60. McGaugh, S. E., Heil, C. S. S., Manzano-Winkler, B., Loewe, L., Goldstein, S., Himmel, T. L., & Noor, M. A. F. (2012). Recombination Modulates How Selection Affects Linked Sites in Drosophila. PLOS Biology, 10(11), e1001422. 10.1371/journal.pbio.1001422

61. McKenna, A., Hanna, M., Banks, E., Sivachenko, A., Cibulskis, K., Kernytsky, A., Garimella, K., Altshuler, D., Gabriel, S., Daly, M., & DePristo, M. A. (2010). The Genome Analysis Toolkit: A MapReduce framework for analyzing next-generation DNA sequencing data. Genome Research, 20(9), 1297–1303. 10.1101/gr.107524.110

62. Miller, D. E., Smith, C. B., Kazemi, N. Y., Cockrell, A. J., Arvanitakis, A. V., Blumenstiel, J. P., Jaspersen, S. L., & Hawley, R. S. (2016). Whole-Genome Analysis of Individual Meiotic Events in Drosophila melanogaster Reveals That Noncrossover Gene Conversions Are Insensitive to Interference and the Centromere Effect. Genetics, 203(1), 159–171. 10.1534/genetics.115.186486

63. Moran, B. M., Payne, C., Langdon, Q., Powell, D. L., Brandvain, Y., & Schumer, M. (2021). The genomic consequences of hybridization. eLife, 10, e69016. 10.7554/eLife.69016

64. Myung, K., Datta, A., Chen, C., & Kolodner, R. D. (2001). SGS1, the Saccharomyces cerevisiae homologue of BLM and WRN, suppresses genome instability and homeologous recombination. Nature Genetics, 27(1), 113–116. 10.1038/83673

65. Nachman, M. & Payseur, B. (2012). Recombination rate variation and speciation: Theoretical predictions and empirical results from rabbits and mice. Philosophical Transactions of the Royal Society B: Biological Sciences, 367(1587), 409–421. 10.1098/rstb.2011.0249

66. Nespolo, R. F., Villarroel, C. A., Oporto, C. I., Tapia, S. M., Vega-Macaya, F., Urbina, K., Chiara, M. D., Mozzachiodi, S., Mikhalev, E., Thompson, D., Larrondo, L. F., Saenz-Agudelo, P., Liti, G., & Cubillos, F. A. (2020). An Out-of-Patagonia migration explains the worldwide diversity and distribution of Saccharomyces eubayanus lineages. PLOS Genetics, 16(5), e1008777. 10.1371/journal.pgen.1008777

67. Petit, M., Astruc, J.-M., Sarry, J., Drouilhet, L., Fabre, S., Moreno, C. R., & Servin, B. (2017). Variation in Recombination Rate and Its Genetic Determinism in Sheep Populations. Genetics, 207(2), 767–784. 10.1534/genetics.117.300123

68. Pool, J. E. (2015). The Mosaic Ancestry of the Drosophila Genetic Reference Panel and the D. melanogaster Reference Genome Reveals a Network of Epistatic Fitness Interactions. Molecular Biology and Evolution, 32(12), 3236–3251. 10.1093/molbev/msv194

69. Raffoux, X., Bourge, M., Dumas, F., Martin, O. C., & Falque, M. (2018). Role of Cis, Trans, and Inbreeding Effects on Meiotic Recombination in Saccharomyces cerevisiae. Genetics, 210(4), 1213–1226. 10.1534/genetics.118.301644

70. Ravinet, M., Yoshida, K., Shigenobu, S., Toyoda, A., Fujiyama, A., & Kitano, J. (2018). The genomic landscape at a late stage of stickleback speciation: High genomic divergence interspersed by small localized regions of introgression. PLOS Genetics, 14(5), e1007358. 10.1371/journal.pgen.1007358

71. Robinson, J. T., Thorvaldsdóttir, H., Winckler, W., Guttman, M., Lander, E. S., Getz, G., & Mesirov, J. P. (2011). Integrative genomics viewer. Nature Biotechnology, 29(1), 24–26. 10.1038/nbt.1754

72. Rockman, M. V., & Kruglyak, L. (2009). Recombinational Landscape and Population Genomics of Caenorhabditis elegans. PLOS Genetics, 5(3), e1000419. 10.1371/journal.pgen.1000419

73. Rogers, D. W., McConnell, E., Ono, J., & Greig, D. (2018). Spore-autonomous fluorescent protein expression identifies meiotic chromosome mis-segregation as the principal cause of hybrid sterility in yeast. PLOS Biology, 16(11), e2005066. 10.1371/journal.pbio.2005066

74. Ruderfer, D. M., Pratt, S. C., Seidel, H. S., & Kruglyak, L. (2006). Population genomic analysis of outcrossing and recombination in yeast. Nature Genetics, 38(9), Article 9. 10.1038/ng1859

75. Samuk, K., Manzano-Winkler, B., Ritz, K. R., & Noor, M. A. F. (2019). *Natural selection shapes variation in genome-wide recombination rate in* Drosophila pseudoobscura [Preprint]. Genetics. 10.1101/787382

76. Sandor, C., Li, W., Coppieters, W., Druet, T., Charlier, C., & Georges, M. (2012). Genetic variants in REC8, RNF212, and PRDM9 influence male recombination in cattle. PLoS Genetics, 8(7), e1002854. 10.1371/journal.pgen.1002854

77. Scannell, D. R., Zill, O. A., Rokas, A., Payen, C., Dunham, M. J., Eisen, M. B., Rine, J., Johnston, M., & Hittinger, C. T. (2011). The Awesome Power of Yeast Evolutionary Genetics: New Genome Sequences and Strain Resources for the Saccharomyces sensu stricto Genus. G3: Genes, Genomes, Genetics, 1(1), 11–25. 10.1534/g3.111.000273

78. Schaeffer, S. W., & Anderson, W. W. (2005). Mechanisms of Genetic Exchange Within the Chromosomal Inversions of Drosophila pseudoobscura. Genetics, 171(4), 1729–1739. 10.1534/genetics.105.041947

79. Schreiber, M., Gao, Y., Koch, N., Fuchs, J., Heckmann, S., Himmelbach, A., Börner, A., Özkan, H., Maurer, A., Stein, N., Mascher, M., & Dreissig, S. (2022). Recombination Landscape Divergence Between Populations is Marked by Larger Low-Recombining Regions in Domesticated Rye. Molecular Biology and Evolution, 39(6), msac131. 10.1093/molbev/msac131

80. Schumer, M., Xu, C., Powell, D. L., Durvasula, A., Skov, L., Holland, C., Blazier, J. C., Sankararaman, S., Andolfatto, P., Rosenthal, G. G., & Przeworski, M. (2018). Natural selection interacts with recombination to shape the evolution of hybrid genomes. Science, 360(6389), 656–660. 10.1126/science.aar3684

81. Sharma, D., Denmat, S. H.-L., Matzke, N. J., Hannan, K., Hannan, R. D., O’Sullivan, J. M., & Ganley, A. R. D. (2022). A new method for determining ribosomal DNA copy number shows differences between *Saccharomyces cerevisiae* populations. Genomics, 114(4), 110430. 10.1016/j.ygeno.2022.110430

82. Shi, J., Wolf, S. E., Burke, J. M., Presting, G. G., Ross-Ibarra, J., & Dawe, R. K. (2010). Widespread Gene Conversion in Centromere Cores. PLoS Biology, 8(3), e1000327. 10.1371/journal.pbio.1000327

83. Singhal, S., Leffler, E. M., Sannareddy, K., Turner, I., Venn, O., Hooper, D. M., Strand, A. I., Li, Q., Raney, B., Balakrishnan, C. N., Griffith, S. C., McVean, G., & Przeworski, M. (2015). Stable recombination hotspots in birds. *Science (New York*, N.Y.), 350(6263), 928–932. 10.1126/science.aad0843

84. Smith, J. M., & Haigh, J. (1974). The hitch-hiking effect of a favourable gene. Genetical Research, 23(1), 23–35.

85. Smukowski, C. S., & Noor, M. a. F. (2011). Recombination rate variation in closely related species. Heredity, 107(6), 496–508. 10.1038/hdy.2011.44

86. Smukowski Heil, C. S., Ellison, C., Dubin, M., & Noor, M. A. F. (2015). Recombining without Hotspots: A Comprehensive Evolutionary Portrait of Recombination in Two Closely Related Species of Drosophila. Genome Biology and Evolution, 7(10), 2829–2842. 10.1093/gbe/evv182

87. Stapley, J., Feulner, P. G. D., Johnston, S. E., Santure, A. W., & Smadja, C. M. (2017). Variation in recombination frequency and distribution across eukaryotes: Patterns and processes. Philosophical Transactions of the Royal Society B: Biological Sciences, 372(1736), 20160455. 10.1098/rstb.2016.0455

88. Stelkens, R., & Bendixsen, D. P. (2022). The evolutionary and ecological potential of yeast hybrids. Current Opinion in Genetics & Development, 76, 101958. 10.1016/j.gde.2022.101958

89. Szymanska-Lejman, M., Dziegielewski, W., Dluzewska, J., Kbiri, N., Bieluszewska, A., Poethig, R. S., & Ziolkowski, P. A. (2023). The effect of DNA polymorphisms and natural variation on crossover hotspot activity in Arabidopsis hybrids. Nature Communications, 14(1), 33. 10.1038/s41467-022-35722-3

90. Talbert, P. B., & Henikoff, S. (2010). Centromeres Convert but Don’t Cross. PLOS Biology, 8(3), e1000326. 10.1371/journal.pbio.1000326

91. Tellini, N., De Chiara, M., Mozzachiodi, S., Tattini, L., Vischioni, C., Naumova, E., Warringer, J., Bergström, A., & Liti, G. (2023). Ancient and recent origins of shared polymorphisms in yeas. 10.21203/rs.3.rs-2573222/v1

92. Tsai, I. J., Burt, A., & Koufopanou, V. (2010). Conservation of recombination hotspots in yeast. Proceedings of the National Academy of Sciences. 10.1073/pnas.0908774107

93. Veller, C., Edelman, N. B., Muralidhar, P., & Nowak, M. A. (2023). Recombination and selection against introgressed DNA. Evolution, 77(4), 1131–1144. 10.1093/evolut/qpad021

94. Veller, C., Kleckner, N., & Nowak, M. A. (2019). A rigorous measure of genome-wide genetic shuffling that takes into account crossover positions and Mendel’s second law. Proceedings of the National Academy of Sciences of the United States of America, 116(5), 1659–1668. 10.1073/pnas.1817482116

95. Wijnker, E., Velikkakam James, G., Ding, J., Becker, F., Klasen, J. R., Rawat, V., Rowan, B. A., de Jong, D. F., de Snoo, C. B., Zapata, L., Huettel, B., de Jong, H., Ossowski, S., Weigel, D., Koornneef, M., Keurentjes, J. J., & Schneeberger, K. (2013). The genomic landscape of meiotic crossovers and gene conversions in Arabidopsis thaliana. eLife, 2, e01426. 10.7554/eLife.01426

96. Zeyl, C. W., & Otto, S. P. (2007). A short history of recombination in yeast. Trends in Ecology & Evolution, 22(5), 223–225. 10.1016/j.tree.2007.02.005

97. Ziolkowski, P. A., Berchowitz, L. E., Lambing, C., Yelina, N. E., Zhao, X., Kelly, K. A., Choi, K., Ziolkowska, L., June, V., Sanchez-Moran, E., Franklin, C., Copenhaver, G. P., & Henderson, I. R. (2015). Juxtaposition of heterozygous and homozygous regions causes reciprocal crossover remodelling via interference during Arabidopsis meiosis. eLife, 4, e03708. 10.7554/eLife.03708

98. Ziolkowski, P. A., & Henderson, I. R. (2017). Interconnections between meiotic recombination and sequence polymorphism in plant genomes. New Phytologist, 213(3), 1022–1029. 10.1111/nph.14265

