## Supplemental Information for "The recombination landscape of introgression in yeast"

### Supplementary Material

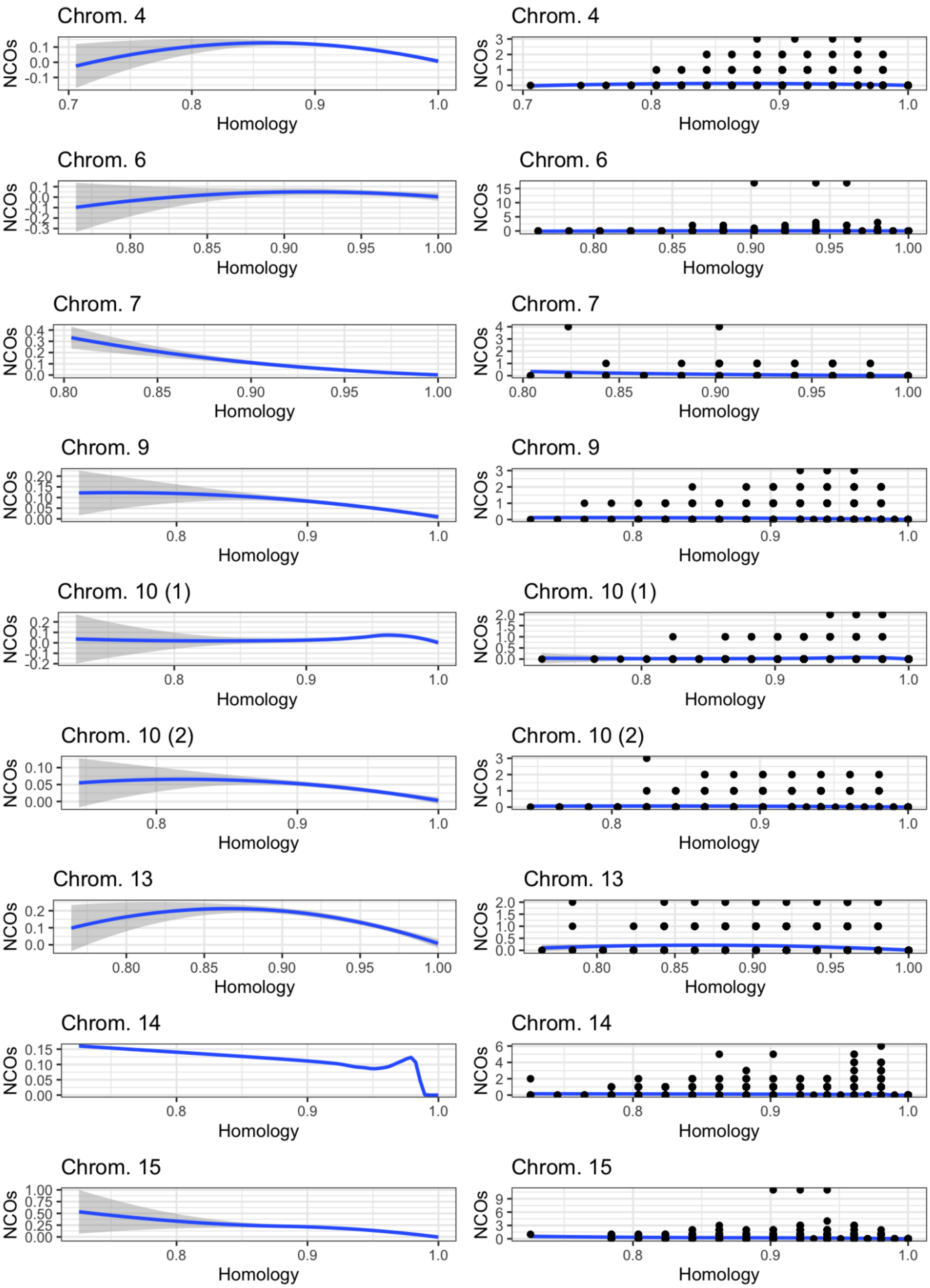

Chrom. 4
NCOs
Homology
Chrom. 4
NCOs
Homology
Chrom. 6
NCOs
Homology
Chrom. 6
NCOs
Homology
Chrom. 7
NCOs
Homology
Chrom. 7
NCOs
Homology
Chrom. 9
NCOs
Homology
Chrom. 9
NCOs
Homology
Chrom. 10 (1)
NCOs
Homology
Chrom. 10 (1)
NCOs
Homology
Chrom. 10 (2)
NCOs
Homology
Chrom. 10 (2)
NCOs
Homology
Chrom. 13
NCOs
Homology
Chrom. 13
NCOs
Homology
Chrom. 14
NCOs
Homology
Chrom. 14
NCOs
Homology
Chrom. 15
NCOs
Homology
Chrom. 15
NCOs
Homology

*Figure S1: Loess regression plots of NCOs as a function of homology in the introgressions of the fermentation cross. The left column's y-axis is scaled to the size of the loess curve, while the right column's y-axis is scaled to the NCO count. The gray shading indicates the standard error for the loess estimates. Chromosome 14's SE ribbon was removed for readability.*

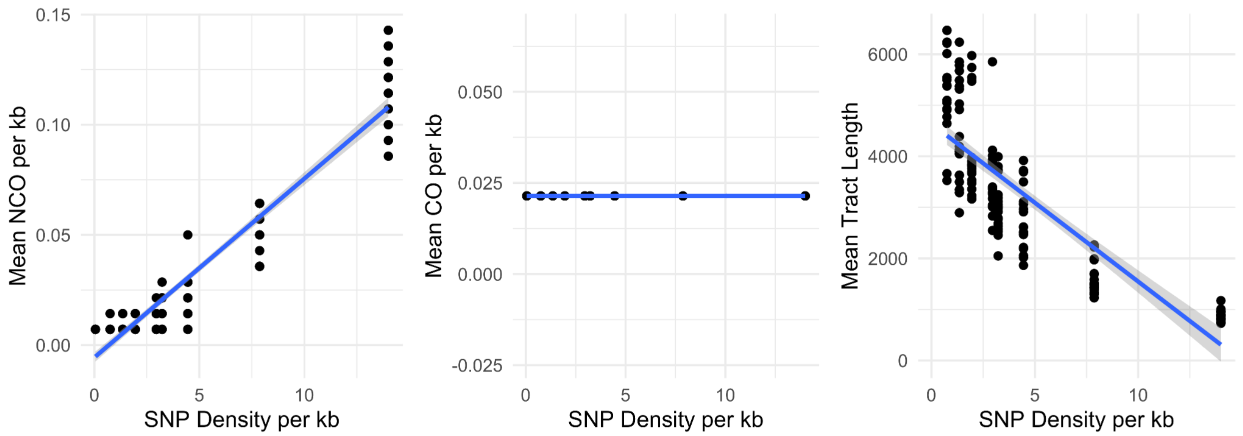


*Figure S2: Downsample densities and NCO rate, CO rate, and tract length. Introgression on chromosome 4.*

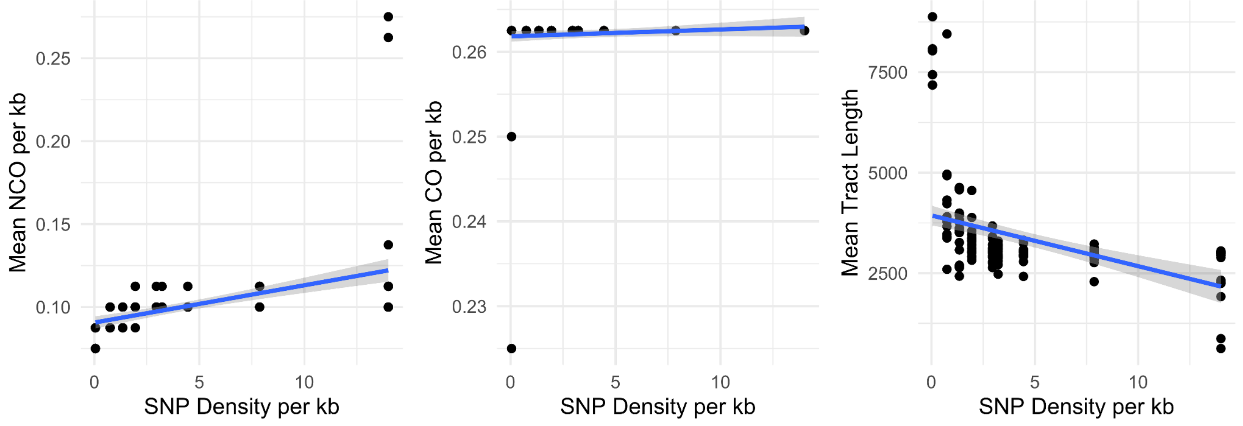


*Figure S3: Downsample densities and NCO rate, CO rate, and tract length. Introgression on chromosome 6.*

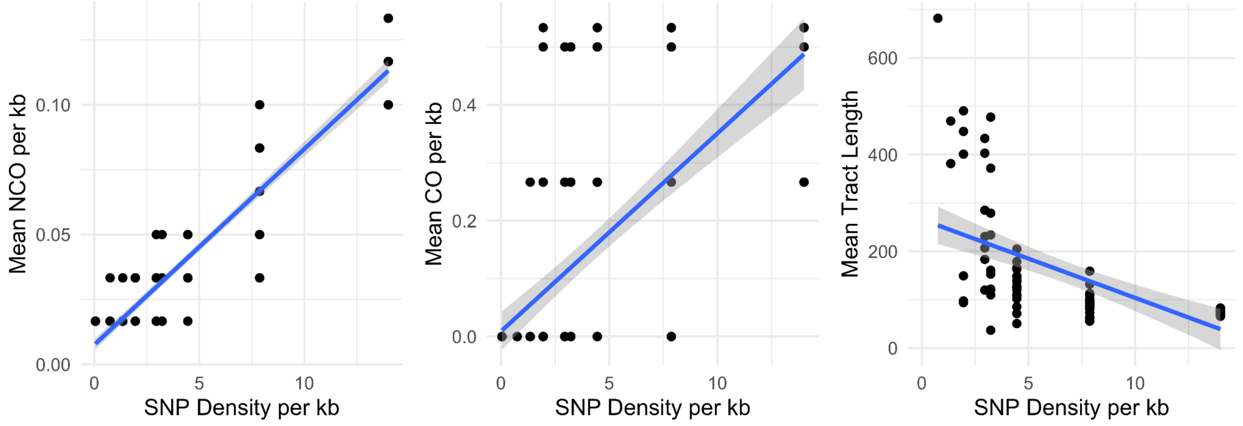

*Figure S4: Downsample densities and NCO rate, CO rate, and tract length. Introgression on chromosome 7.*

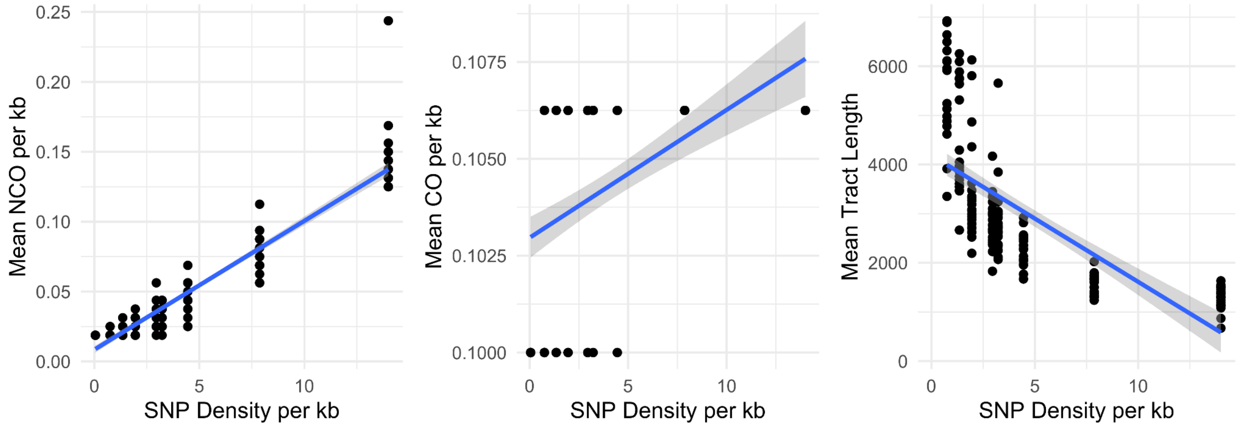


*Figure S5: Downsample densities and NCO rate, CO rate, and tract length. Introgression on chromosome 9.*

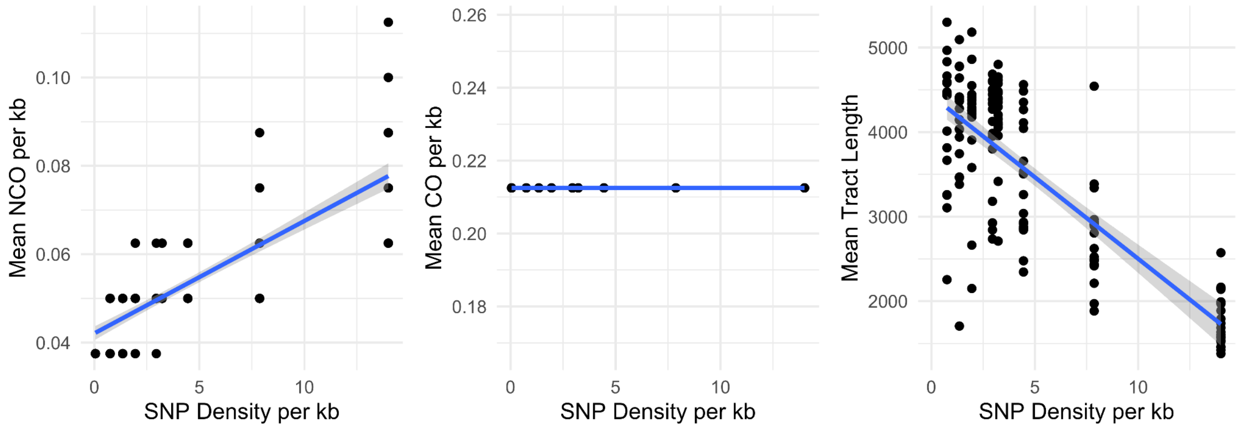


*Figure S6: Downsample densities and NCO rate, CO rate, and tract length. First introgression on chromosome 10 (10a).*

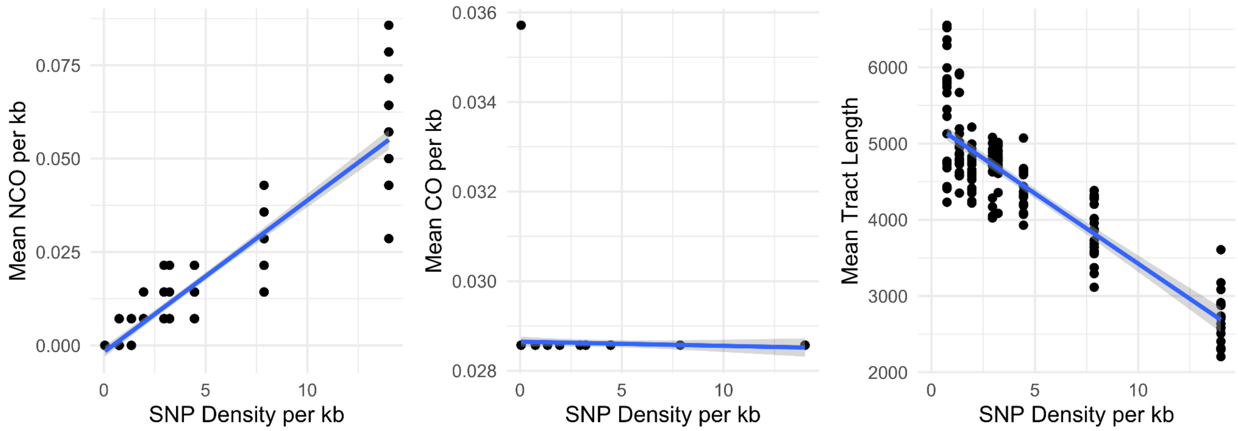


*Figure S7: Downsample densities and NCO rate, CO rate, and tract length. First introgression on chromosome 10 (10b).*

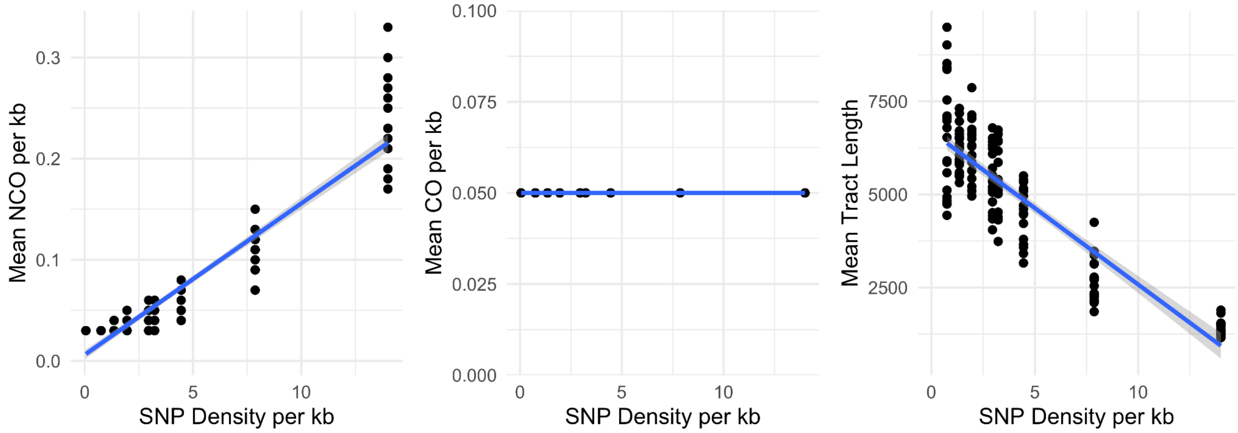


*Figure S8: Downsample densities and NCO rate, CO rate, and tract length. Introgression on chromosome 13.*

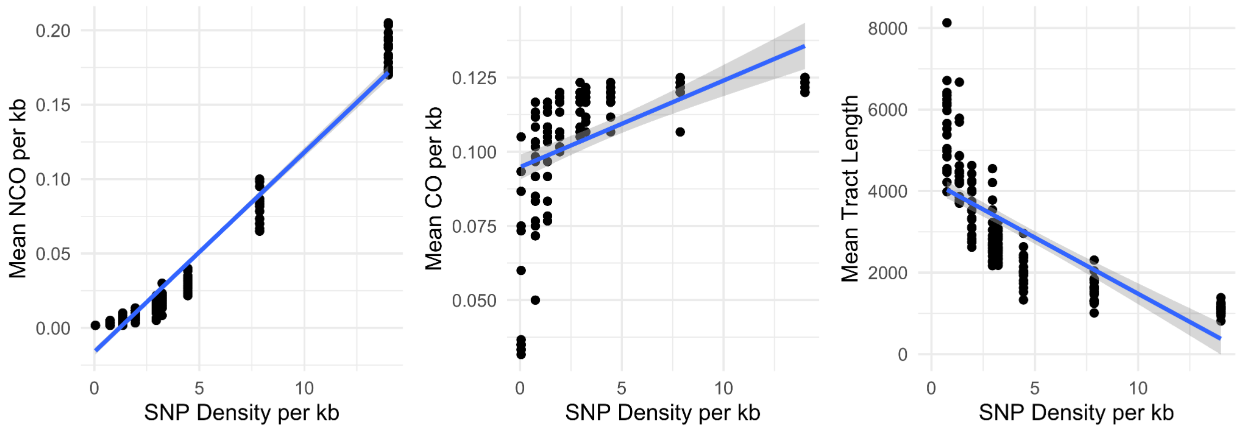

*Figure S9: Downsample densities and NCO rate, CO rate, and tract length. Introgression on chromosome 14.*

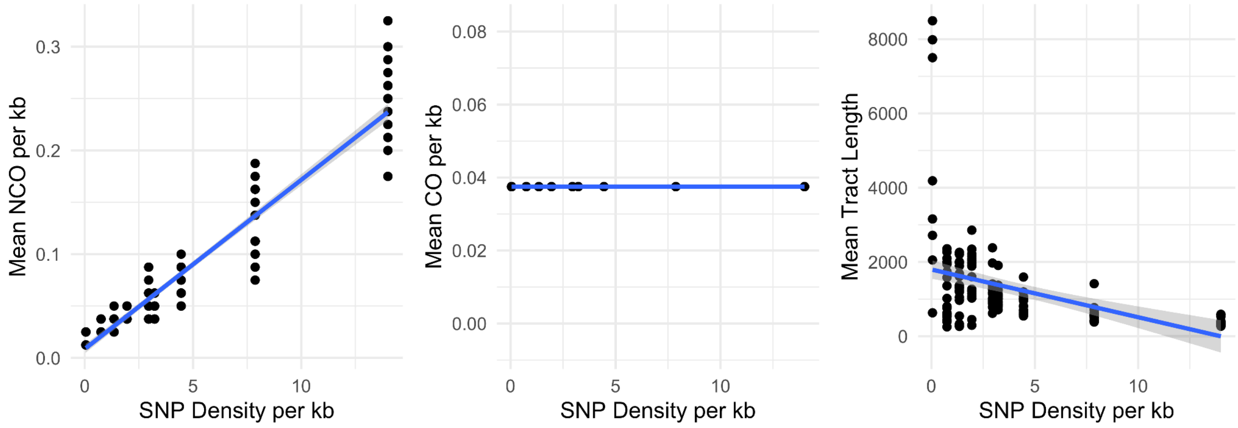


*Figure S10: Downsample densities and NCO rate, CO rate, and tract length. Introgression on chromosome 15.*

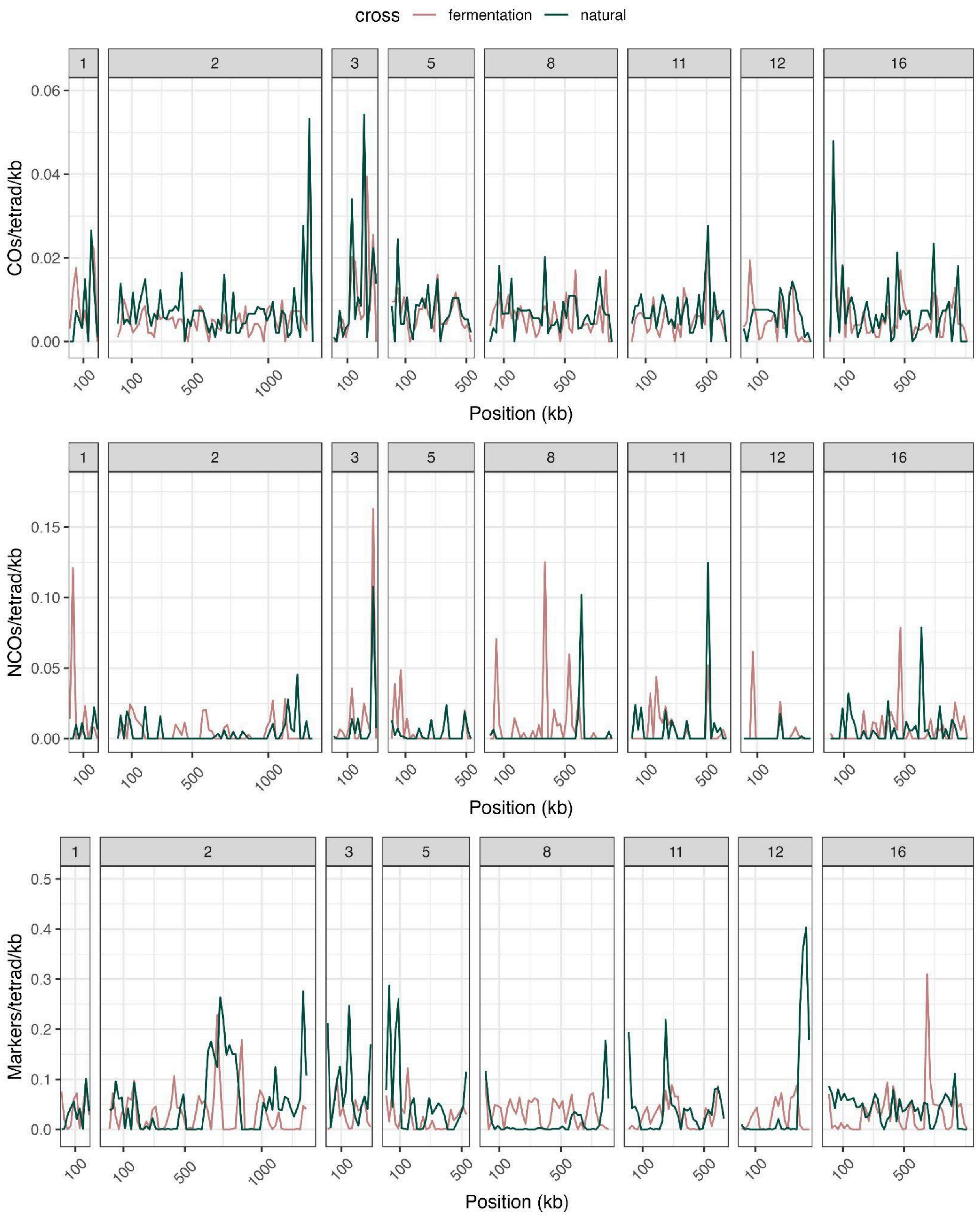


*Figure S11:* S. uvarum *chromosomes not containing introgressions split into 20kb, non-overlapping windows. CO, NCO, and SNP counts are reported for both crosses*

*(fermentation and natural). CO counts are smoothed when the true location of the CO split could be in one of multiple windows. NCO counts are corrected for marker resolution.*

*Table S1: Strain information*

| Strain number | Strain name | Genotype | Isolation source | Isolation location | Cross | NCBI SRA | Citation | Obtained from |
|---|---|---|---|---|---|---|---|---|
| yCSH347 | yHCT78 | | Bark of Quercus acutissima | Chaumette Vineyard, Ste. Genevieve, Missouri | | SRR1119189 | Almeida et al 2014 | Chris Hittinger |
| yCSH345 | UCD61-137 | | Drosophila pseudoobscura | Berryessa Hills, California | | SRR1119180 | Almeida et al 2014 | Portugese Yeast Culture Collection (PYCC 6878) |
| yCSH561 | DBVPG 7787 | | Wine | Slovakia | | SRR1119199 | Almeida et al 2014 | Portugese Yeast Culture Collection (PYCC6876) |
| yCSH35 | GM14 | | Fermenting grape must | France | | SRR1119200 | Almeida et al 2014 | Portugese Yeast Culture Collection (PYCC 6892) |
| yCSH833 | GM14 | HOdelta0::kanMX MATa | | | European fermentation | | This study | |
| yCSH836 | DBVPG 7787 | HOdelta0::kanMX MATx | | | European fermentation | | This study | |
| yCSH837 | UCD61-137 | HOdelta0::kanMX MATa | | | North American natural | | This study | |
| yCSH840 | yHCT78 | HOdelta0::kanMX MATx | | | North American natural | | This study | |

| Plasmid strain number | | |
|---|---|---|
| pCSH2 | | pFA6a-TEF2Pr-dTomato-ADH1-Primer-KANMX6 |

*Table S2: Copy number variation in GM14*

| Chromosome | Start | End | Size | Type |
|---|---|---|---|---|
| 5 | 294582 | 306485 | 11903 | amplification |
| 7 | 840000 | 859500 | 19500 | amplification |
| 15 | 414372 | 416662 | 2290 | amplification |

*Table S3: Location of introgressions*

| Chromosome | Start Position | End position |
|---|---|---|
| 4 | 866500 | 983774 |
| 6 | 1 | 65500 |
| 7 | 1 | 53500 |
| 9 | 158500 | 298500 |
| 10 | 234500 | 288500 |
| 10 | 301500 | 428500 |
| 13 | 26500 | 103500 |
| 14 | 18500 | 586500 |
| 15 | 367500 | 434500 |

*Table S4: Mean and standard error of CO counts per chromosome for fermentation and natural crosses.*

| Chromosome | Natural count | Natural SE | Fermentation count | Fermentation SE |
|---|---|---|---|---|
| 1 | 1.3958 | 0.1254 | 1.9574 | 0.0960 |
| 2 | 9.2083 | 0.3150 | 6.8298 | 0.2136 |
| 3 | 3.5000 | 0.1786 | 2.9362 | 0.1503 |
| 4 | 7.2292 | 0.2650 | 4.5319 | 0.1796 |
| 5 | 3.8750 | 0.1921 | 3.5106 | 0.1517 |
| 6 | 4.0208 | 0.1412 | 2.8936 | 0.1753 |
| 7 | 6.0208 | 0.2218 | 5.7234 | 0.2389 |
| 8 | 5.3750 | 0.2834 | 4.7021 | 0.1548 |
| 9 | 2.8333 | 0.2005 | 2.0638 | 0.1533 |
| 10 | 8.1667 | 0.2798 | 5.0638 | 0.2460 |
| 11 | 4.5208 | 0.2041 | 3.5319 | 0.1211 |
| 12 | 2.8958 | 0.1272 | 2.2340 | 0.1557 |
| 13 | 6.0625 | 0.2371 | 4.5957 | 0.1891 |
| 14 | 5.4167 | 0.2576 | 2.8936 | 0.1881 |
| 15 | 5.4583 | 0.2018 | 4.3404 | 0.2069 |
| 16 | 6.5625 | 0.2695 | 5.8511 | 0.2040 |

*Table S5: Mean and standard error of NCO counts per chromosome for fermentation and natural crosses.*

| Chromosome | Natural count | Natural SE | Fermentation count | Fermentation SE |
|---|---|---|---|---|
| 1 | 0.3958 | 0.1020 | 0.7447 | 0.1376 |
| 2 | 2.3958 | 0.2340 | 1.7872 | 0.2948 |
| 3 | 1.5208 | 0.1657 | 0.6809 | 0.1294 |
| 4 | 2.2292 | 0.2442 | 3.1277 | 1.2153 |
| 5 | 1.0625 | 0.1746 | 0.4894 | 0.1249 |
| 6 | 1.8750 | 0.1944 | 1.4468 | 0.1943 |
| 7 | 2.2083 | 0.2325 | 2.1915 | 0.3658 |
| 8 | 0.5000 | 0.1074 | 1.4043 | 0.1891 |
| 9 | 1.3333 | 0.1745 | 2.1702 | 0.8392 |
| 10 | 2.6250 | 0.2787 | 3.2766 | 1.0574 |
| 11 | 1.3333 | 0.1719 | 1.1064 | 0.1616 |
| 12 | 0.2083 | 0.0592 | 0.5745 | 0.1451 |
| 13 | 2.4375 | 0.2695 | 3.3617 | 1.2270 |
| 14 | 1.8750 | 0.1921 | 7.9149 | 4.0521 |
| 15 | 1.5417 | 0.1929 | 3.2340 | 1.0061 |
| 16 | 1.9375 | 0.2256 | 1.3404 | 0.2001 |

*Table S6: Spearman's correlations between crosses in non-introgressed regions*

| Chrom | CO correlation | CO correlation p-value | NCO correlation | NCO correlation p-value | SNP correlation | SNP correlation p-value |
|---|---|---|---|---|---|---|
| 1 | 0.6667 | 0.0353 | –0.3311 | 0.3500 | -0.0061 | 0.9867 |
| 2 | 0.2302 | 0.0650 | 0.0069 | 0.9564 | 0.0086 | 0.9459 |
| 3 | 0.3816 | 0.1604 | 0.0120 | 0.9661 | -0.3339 | 0.2238 |
| 4 | 0.2205 | 0.1553 | 0.2558 | 0.0978 | 0.3420 | 0.0248 |
| 5 | 0.5205 | 0.0054 | 0.2386 | 0.2308 | -0.0746 | 0.7117 |
| 6 | 0.5908 | 0.0030 | -0.2015 | 0.3566 | 0.2370 | 0.2762 |
| 7 | 0.2048 | 0.1580 | 0.1016 | 0.4871 | 0.0106 | 0.9425 |
| 8 | 0.2187 | 0.1695 | -0.1107 | 0.4908 | -0.2473 | 0.1190 |
| 9 | 0.1278 | 0.6774 | 0.0635 | 0.8366 | -0.5992 | 0.0305 |
| 10 | 0.2313 | 0.1510 | 0.2300 | 0.1533 | 0.2353 | 0.1438 |
| 11 | 0.5733 | 0.0006 | 0.2645 | 0.1436 | 0.2297 | 0.2060 |
| 12 | 0.4717 | 0.0231 | 0.2589 | 0.2329 | -0.2188 | 0.3157 |
| 13 | 0.2825 | 0.0735 | -0.1327 | 0.4081 | 0.1341 | 0.4032 |
| 14 | 0.8932 | 0.0028 | -0.2744 | 0.5107 | -0.6831 | 0.0618 |
| 15 | 0.4660 | 0.0048 | 0.3021 | 0.0778 | 0.3243 | 0.0573 |
| 16 | 0.4318 | 0.0027 | -0.0689 | 0.6492 | -0.0898 | 0.5528 |

*Table S7: Spearman's correlations between crosses in introgressed regions*

| Chrom. | CO correlation | CO correlation p-value | NCO correlation | NCO correlation p-value | SNP correlation | SNP correlation p-value |
|---|---|---|---|---|---|---|
| 4 | 0.2118 | 0.6484 | 0.2883 | 0.5307 | 0.7143 | 0.0881 |
| 6 | 0.4000 | 0.7500 | 0.7746 | 0.2254 | -0.4000 | 0.7500 |
| 7 | 0.0000 | 1.0000 | NA | NA | -0.8660 | 0.3333 |
| 9 | -0.2547 | 0.5427 | 0.6587 | 0.0757 | 0.1928 | 0.6474 |
| 10 | -0.2071 | 0.5411 | 0.2642 | 0.4324 | 0.2727 | 0.4182 |
| 13 | 0.0000 | 1.0000 | NA | NA | 0.1539 | 0.8048 |
| 14 | 0.2274 | 0.2269 | -0.0416 | 0.8274 | -0.3782 | 0.0393 |
| 15 | 0.2000 | 0.9167 | -0.8000 | 0.3333 | 0.8000 | 0.3333 |

*Table S8: $\bar{r}$ values for whole chromosomes*

| Chromosome | Natural mean | Natural SE | Fermentation mean | Fermentation SE | t-test p-value |
|---|---|---|---|---|---|
| 1 | 0.2615 | 0.0140 | 0.2651 | 0.0132 | 0.8498 |
| 2 | 0.4264 | 0.0068 | 0.3993 | 0.0090 | 0.0172 |
| 3 | 0.3101 | 0.0121 | 0.3104 | 0.0131 | 0.9848 |
| 4 | 0.4166 | 0.0067 | 0.3728 | 0.0100 | 0.0003 |
| 5 | 0.3606 | 0.0106 | 0.3267 | 0.0125 | 0.0391 |
| 6 | 0.3672 | 0.0106 | 0.3265 | 0.0130 | 0.0158 |
| 7 | 0.4034 | 0.0085 | 0.3956 | 0.0091 | 0.5307 |
| 8 | 0.3680 | 0.0108 | 0.3803 | 0.0096 | 0.3968 |
| 9 | 0.3124 | 0.0133 | 0.2269 | 0.0135 | $8.75 \times 10^{-6}$ |
| 10 | 0.4061 | 0.0074 | 0.3655 | 0.0105 | 0.0018 |
| 11 | 0.3674 | 0.0103 | 0.3388 | 0.0114 | 0.0634 |
| 12 | 0.3583 | 0.0115 | 0.2699 | 0.0141 | $1.9 \times 10^{-6}$ |
| 13 | 0.3784 | 0.0098 | 0.3896 | 0.0099 | 0.4228 |
| 14 | 0.3758 | 0.0106 | 0.2163 | 0.0114 | $<2.2 \times 10^{-16}$ |
| 15 | 0.4064 | 0.0076 | 0.3582 | 0.0112 | 0.0004 |
| 16 | 0.3991 | 0.0089 | 0.3808 | 0.0103 | 0.1794 |

*Table S9: $\bar{r}$ values for introgressed regions*

| Chromosome | Natural mean | Natural SE | Fermentation mean | Fermentation SE | t-test p-value |
|---|---|---|---|---|---|
| 4 | 0.1539 | 0.0131 | 0.0161 | 0.0059 | $<2.2 \times 10^{-16}$ |
| 6 | 0.0577 | 0.0111 | 0.0175 | 0.0058 | 0.0015 |
| 7 | 0.2678 | 0.0075 | 0.1551 | 0.0155 | $2.78 \times 10^{-10}$ |
| 9 | 0.1826 | 0.0146 | 0.0393 | 0.0087 | $1.15 \times 10^{-15}$ |
| 10a | 0.0604 | 0.0106 | 0.0123 | 0.0043 | $3.46 \times 10^{-5}$ |
| 10b | 0.2017 | 0.0152 | 0.0075 | 0.0029 | $<2.2 \times 10^{-16}$ |
| 13 | 0.1040 | 0.0140 | 0.0193 | 0.0055 | $4.65 \times 10^{-8}$ |
| 14 | 0.3364 | 0.0126 | 0.1513 | 0.0120 | $<2.2 \times 10^{-16}$ |
| 15 | 0.1596 | 0.0153 | 0.0242 | 0.0056 | $6.81 \times 10^{-15}$ |

*Table S10: Summaries of linear regression for downsampled introgressions. Each regression has either CO count, NCO count or NCO tract length as the response variable, and marker density as the independent variable.*

| Chromosome | Response Variable | Intercept | Intercept p-value | Marker dens | Marker dens p-value | Adjusted R-squared |
|---|---|---|---|---|---|---|
| 4 | CO | 0.4286 | 0 | 0.0000 | 0.3334 | 0.5002 |
| 4 | NCO | -0.1122 | 0 | 0.1623 | 0 | 0.9003 |
| 4 | Tract lenght | 4636.396 | 0 | 309.0741 | 0 | 0.6948 |
| 6 | CO | 5.2364 | 0 | 0.0016 | 0 | 0.2146 |
| 6 | NCO | 1.8130 | 0 | 0.0450 | 0.1426 | 0.0120 |
| 6 | Tract length | 3935.625 | 0 | -126.2340 | 0 | 0.2002 |
| 7 | CO | 0.1782 | 0.5991 | 0.6840 | 0 | 0.4360 |
| 7 | NCO | 0.1516 | 0 | 0.1508 | 0 | 0.8902 |
| 7 | Tract length | 265.9691 | 0 | -16.2048 | 0 | 0.3279 |
| 9 | CO | 2.0593 | 0 | 0.0066 | 0 | 0.2229 |
| 9 | NCO | 0.1689 | 0 | 0.1842 | 0 | 0.9053 |
| 9 | Tract length | 4187.5 | 0 | -257.4853 | 0 | 0.5112 |
| 10a | CO | 4.25 | 0 | 0 | 0.3334 | 0.4991 |
| 10a | NCO | 0.8418 | 0 | 0.0509 | 0 | 0.6643 |
| 10a | Tract length | 4432.434 | 0 | -193.1295 | 0 | 0.6062 |
| 10b | CO | 0.5730 | 0 | -0.0002 | 0.3334 | 0.0053 |
| 10b | NCO | -0.0357 | 0.0161 | 0.0812 | 0 | 0.8526 |
| 10b | Tract length | 5268.21 | 0 | -184.3647 | 0 | 0.7689 |
| 13 | CO | 1 | 0 | 0 | 0.3334 | 0.4993 |
| 13 | NCO | 0.1219 | 0.01 | 0.2993 | 0 | 0.8854 |
| 13 | Tract length | 6682.40 | 0 | -410.3597 | 0 | 0.7844 |
| 14 | CO | 1.8966 | 0 | 0.0583 | 0 | 0.2614 |
| 14 | NCO | -0.3230 | 0 | 0.2683 | 0 | 0.9562 |

| | | | | | | |
|---|---|---|---|---|---|---|
| 14 | Tract length | 4239.71 | 0 | -276.0607 | 0 | 0.5724 |
| 15 | CO | 0.750 | 0 | 0 | 0.3334 | 0.5009 |
| 15 | NCO | 0.173 | 0.0004 | 0.3260 | 0 | 0.9130 |
| 15 | Tract length | 1796.619 | 0 | -128.5568 | 0 | 0.1834 |
